# Artificial environment impact: O2 concentration changes between IVM and IVF alter embryo production, metabolism, and epigenetic marks

**DOI:** 10.1101/2023.11.28.569089

**Authors:** Jessica B Cruz, Carolina M Nogueira, Juliano R Sangalli, Ricardo P Nociti, Dewison R Ambrizi, Alessandra Bridi, Jorge Pinzon, Maira BR Alves, Vera FMH de Lima, Yeda F Watanabe, Fabiana F Bressan, Flavio V Meirelles, Rafael V Sampaio

**Affiliations:** Department of Veterinary Medicine, Faculty of Animal Sciences and Food Engineering, University of Sao Paulo, Pirassununga, Sao Paulo, Brazil; São Paulo State University (UNESP), Jaboticabal, São Paulo, Brazil; Vitrogen – Biotecnologia em Reprodução Animal, Cravinhos, SP, Brazil; STgenetics, Navasota, TX, 77868, USA

**Author notes:** **Corresponding author**. STgenetics, 22575 Hwy 6, Navasota, TX, USA, **E-mail address**. These authors contributed equally to this work (Jessica B Cruz, Carolina M Nogueira).

**Keywords:** epigenetics, cattle, development, culture

## Abstract

Creating an optimal in vitro cell culture environment requires careful simulation of all critical components, especially the gaseous atmosphere. Although it is well-documented that embryonic culture under low oxygen tension promotes embryonic development, little is known about the effect of changes in in vitro maturation (IVM) and fertilization (IVF) on the epigenome. This study explores the role of oxygen tension variation in the early stages of in vitro production of bovine embryos and its impact on oxidative stress and epigenetic remodeling. We initially validated our system by scrutinizing the epigenetic effects on bovine fibroblasts. We observed that cell cultures under 20% O2 exhibited reduced H3K9me2 levels in early passages, which stabilized with prolonged cultivation and elevated gene expression of HIF2a and KDM5C. Our results reveal that oocytes maturing in 20% O2 environments have heightened levels of reactive oxygen species (ROS) and glutathione (GSH), whereas blastocyst embryos maturing under reduced oxygen tension exhibit increased oxidative stress markers (*NRF2*, *SOD1*, *SOD2*), with upregulated transcripts observed in epigenetic remodelers (*KDM5A*, *TET1*). Elevated O2 levels in both IVM and IVF processes showcase improved embryo production. Maintaining consistent O2 levels at either 5% or 20% between IVM and IVF results in heightened GSH, reduced ROS, and increased levels of H3K9me2/3 in embryos. Finally, distinct DNA methylation patterns emerge, indicating higher levels in groups matured under low O2 tension and increased DNA hydroxymethylation in groups fertilized under low O2 tension. In conclusion, our comprehensive investigation underscores the critical role of oxygen concentration in shaping the epigenetic landscape during the early stages of in vitro culture. These findings provide valuable insights for optimizing conditions in assisted reproductive technologies.

## INTRODUCTION

Eukaryotic cellular respiration is fundamentally oxygen-dependent. Although atmospheric oxygen comprises 21%, most mammalian tissues experience lower concentrations, ranging from 3% to 10% (Hancock, Dunne, Walport, Flashman, & Kawamura, 2015). During initial embryonic development in ideal in vivo conditions, oxygen levels fluctuate between 5% and 7% (Fischer & Bavister, 1993; A. J. Harvey et al., 2007). Despite advancements in vitro embryo production (IVP), factors like medium composition, temperature, pH, and oxygen tension in the artificial environment can negatively influence mammalian embryonic development (Menezo, Silvestris, Dale, & Elder, 2016). Incubation conditions in IVP systems may induce oxidative stress, impacting oocytes and inhibiting in vitro embryonic development (Nabenishi et al., 2012; Oyamada & Fukui, 2004).

The link between epigenetic modifications and oxygen concentration is evident through dioxygenase proteins, a major group responsible for epigenetic marks, relying on α-ketoglutarate (αKG) and Fe (II), necessitating oxygen as a substrate (van der Knaap & Verrijzer, 2016). Lysine demethylases (KDMs) and DNA demethylases (TET proteins) are affected by oxygen levels. Hypoxia induces heterodimerization with hypoxia-inducible transcription factor (HIF), influencing intracellular pathways (Loenarz & Schofield, 2008; Semenza, 2002). HIF-1, crucial in cellular events like embryonic development and apoptosis, regulates genes in adaptive pathways (Semenza, 2003). HIF binds DNA sequences, modifying gene transcription. Enzymes for post-translational histone modifications, including methylation, acetylation, phosphorylation, and DNA methylation/demethylation, use oxygen as a substrate, impacting genetic remodeling with varying oxygen concentrations (Martinez & Hausinger, 2015; van der Knaap & Verrijzer, 2016).

Assisted reproduction techniques face challenges linked to their sensitivity to epigenetic remodeling changes, a multi-step process (Ross & Sampaio, 2018). Modulating embryo culture conditions across different oxygen tensions elicits changes in DNA methylation patterns (Bomfim et al., 2017) and histone modifications (Gaspar et al., 2015), thereby influencing imprinted gene regulation in embryonic stem cells (Skiles et al., 2018). Given the pivotal role and intricate influence of oxygen on embryonic development, diverse strategies have emerged to counter its adverse effects, encompassing oxygen reduction to 5% or even ultra-low levels at 2%, as well as the use of antioxidants and endoplasmic reticulum stress inhibitors (Marsico et al., 2023). However, a notable research gap exists in investigating the specific impact of oxygen tension during maturation and in vitro fertilization and its consequential influence on epigenetics—a crucial aspect profoundly impacting the success of ARTs.

This research examines how different oxygen levels during cell culture, maturation, and fertilization affect the expression of genes related to oxidative stress and epigenetic processes in fibroblasts, oocytes, cumulus cells, and bovine embryos. Our study reveals the impact of oxidative stress on metabolism, global DNA methylation, and histone marks in embryos and cells.

## RESULTS

### Oxygen-Modulated Gene Expression and H3K9me2 Levels in Cultured Bovine Fibroblasts

We cultured seven skin-derived fibroblast cell lines over extended periods (P5, P10, and P20), subjecting them to two oxygen atmospheres: high tension (20%) resembling typical incubator condition and low tension (5%), mirroring physiological levels. Gene expression analysis of target genes, including *TET1*, *TET2*, *TET3*, *KDM3A*, *KDM3B*, *KDM4A*, *KDM4C*, *KDM5A*, *KDM5B*, *KDM5C*, *KDM6A*, *KDM6B*, *HIF1a*, *HIF1B*, *HIF2a*, and *HIF3B*, revealed statistical differences in *HIF2a* and *KDM5C* gene expression at the fifth passage (Supplementary Figure S1) (Supplementary Table), indicating cellular heterogeneity and potential adaptation. However, this distinction diminished in prolonged cultures under high and low O2 tensions. In the fifth passage, Immunofluorescence analysis revealed lower levels of H3K9me2 at 20% compared to 5% oxygen tension (Figure 1 A, B). However, in the twenty-second passage, the levels stabilized and were similar between the two oxygen tensions (Figure 1 C, D). The gene expression analysis showed no changes (Supplementary Figure S2).

**Figure 1.**
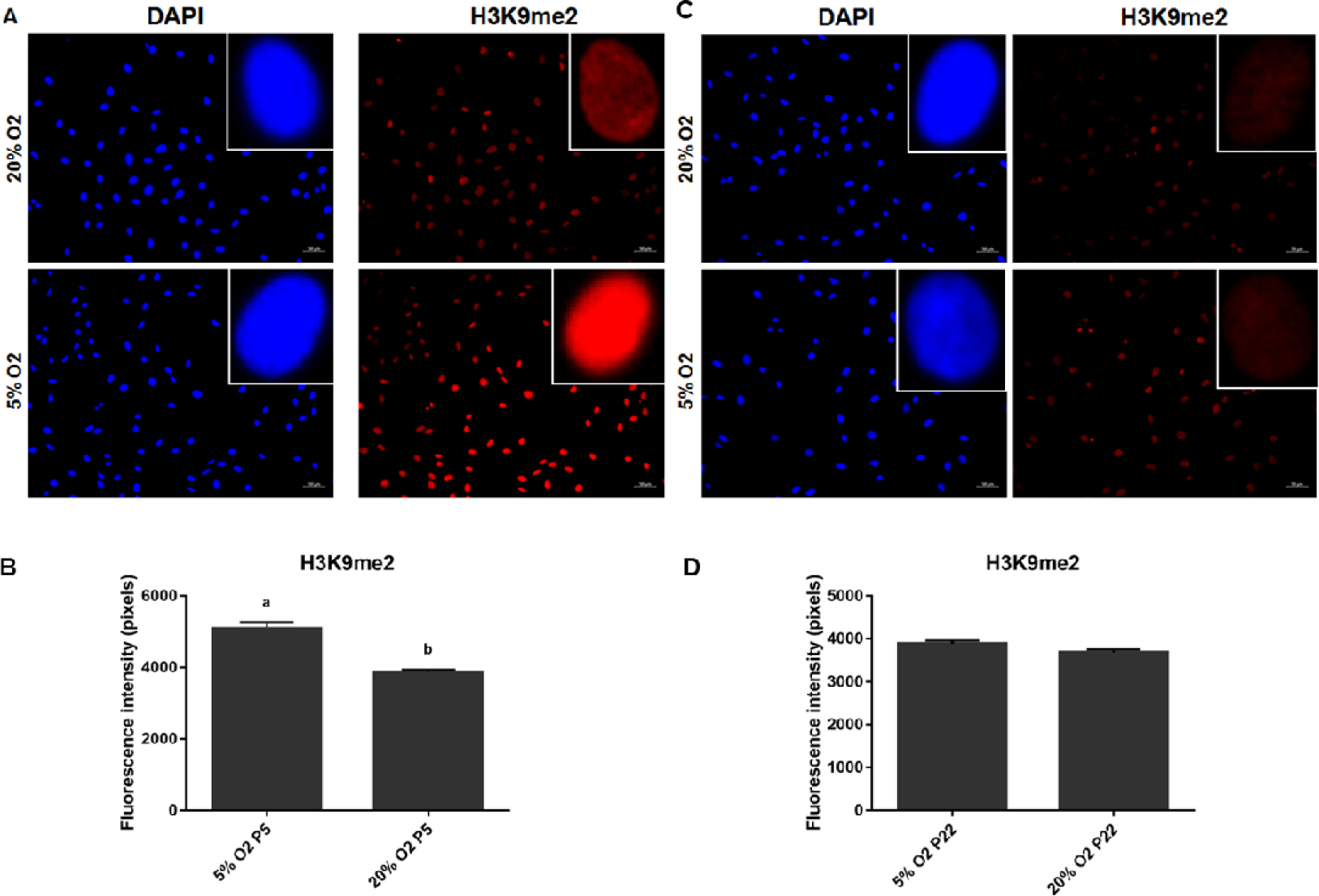
Global levels of H3K9me2 in bovine fibroblasts cultured under different oxygen tensions (5% and 20%) for five and 22 passages. (**A**) Photomicrograph of fibroblasts stained with anti-H3K9me2 antibody (red) and Hoechst (blue) at passage 5. (**B**) Quantification of H3K9me2 levels through nuclear measurements of bovine fibroblasts at passage 5. (**C**) Photomicrograph of fibroblasts stained with anti-H3K9me2 antibody (red) and Hoechst (blue) at passage 22. (**D**) Quantification of H3K9me2 levels through nuclear measurements of bovine fibroblasts at passage 22. Magnification of 200x in epifluorescence microscopy. Isolated bovine fibroblast nucleus magnified 21x compared to others. Student’s t-test revealed statistical significance for the H3K9me2 mark at passage 5 (p < 0.0001).

### High Oxygen concentration during maturation induces ROS and GSH production

We then investigated the effects of different oxygen tensions on oocyte maturation and gene expression. Firstly, we examined the nuclear maturation rate in oocytes exposed to high (M20) and low (M5) oxygen levels. Our findings showed no significant difference in maturation rates between the two groups (p=0.1639). Next, we analyzed the levels of ROS and GSH in oocytes that were matured under high and low oxygen levels using fluorescent probes (Figure 2A). We observed that oocytes exposed to 20% oxygen exhibited higher levels of ROS and GSH (Figure 2B). Further, we investigated the expression patterns of genes related to oxidative stress and epigenetic remodeling in oocytes and cumulus cells. Our comprehensive gene expression analysis demonstrated no significant changes in the expression patterns of genes related to oxidative stress and epigenetic remodeling in oocytes following exposure to different oxygen levels during in vitro maturation (Supplementary Figure S3). Interestingly, we observed elevated transcript levels of the *SOD1* gene in cumulus cells, a key player in the cellular oxidative stress response pathway, with higher expression under high oxygen tension than low oxygen levels (Supplemental Figure S4).

**Figure 2.**
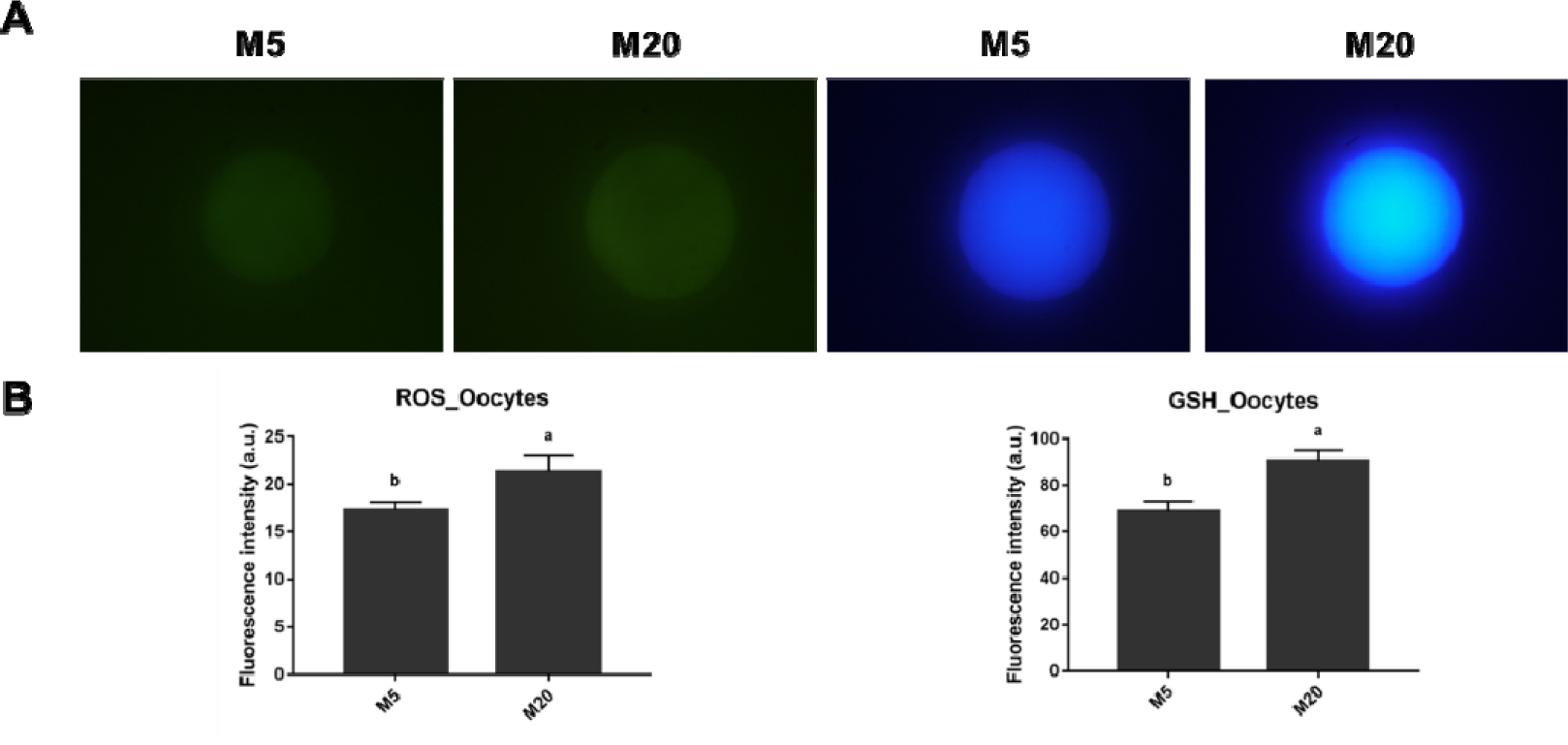
Quantification of intracellular levels of reactive oxygen species (in arbitrary units of fluorescence) in oocytes matured under low (M5) and high (M20) oxygen tension. (A) Epifluorescence photomicrographs of oocytes labeled with a probe for detecting reactive oxygen species (ROS). For the oocyte ROS quantification, the fluorescence intensity of the CellROX Green dye was measured on a Zeis epifluorescence microscope using a 40X objective and Epifluorescence photomicrographs of oocytes labeled with a probe for GSH detection. For the quantification of oocyte GSH, the fluorescence intensity of the Celltracker Blue CMF2HC dye was measured on a Zeiss epifluorescence microscope using a 40X objective. (B)The bars in the graph represent the means, and the error bars represent the standard error of the mean. Different letters (a, b) on the bars indicate a significant difference (p<0.05) by the Kruskal-Wallis test.

### Impact of Oxygen Concentration on Fertilization Rates and In Vitro Production of Bovine Embryos

Our investigation aimed to assess the impact of varying O2 concentrations on fertilization rates, categorizing presumptive zygotes as non-fertilized, polyspermic, or normally fertilized and comparing their respective percentages (Table 1, Supplemental Figure S5). Our findings demonstrated no significant differences in total (p=0.9207), normal (p=0.9880), or polyspermic fertilization (p=0.8585) among different oxygen tensions during maturation and fertilization. Similarly, oxygen levels during in vitro maturation (IVM) and in vitro fertilization (IVF) showed no significant differences in total fertilization rate (IVM (p=0.2718) and IVF (p=0.7633)), normal fertilization rate (IVM (p=0.5375) and IVF (p=0.6970)), or polyspermy (IVM (p=0.3389) and IVF (p=0.2825)) (Table1).

**Table 1.**
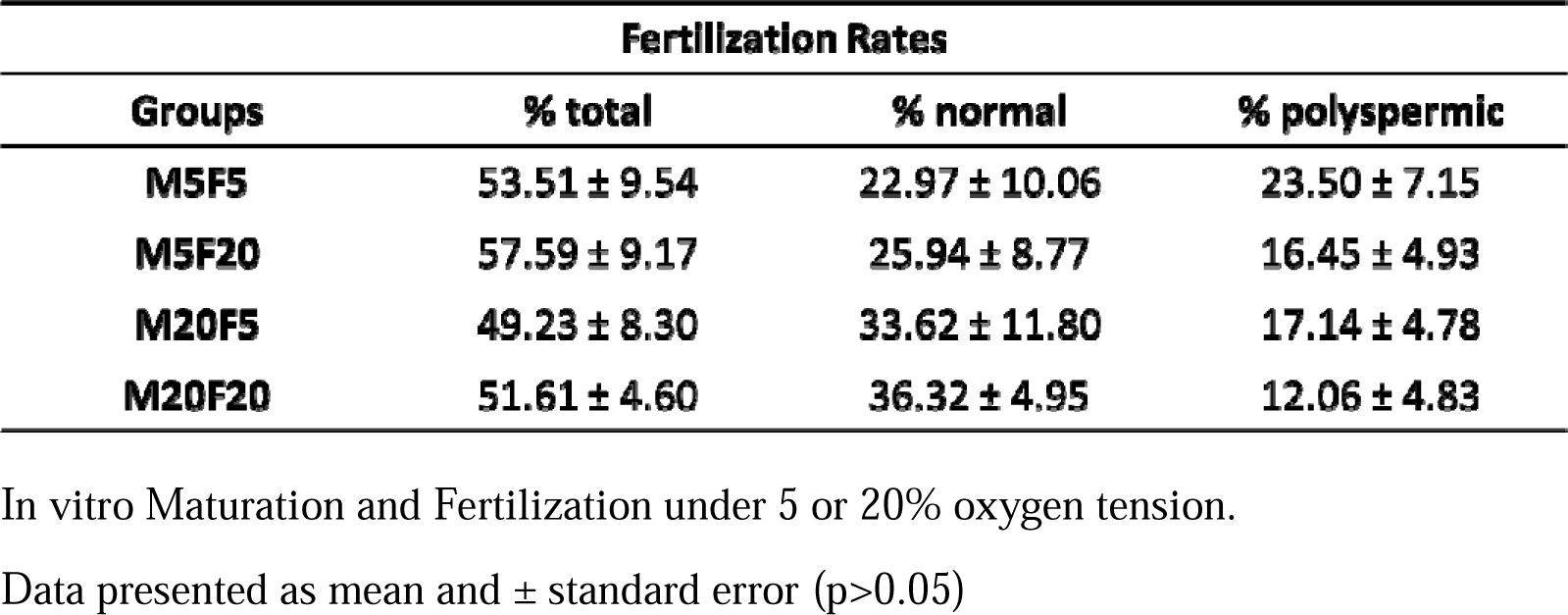
Effect of different O2 tension during IVF on fertilization rates.

| **Fertilization Rates** |  |  |  |
| --- | --- | --- | --- |
| **Groups** | **% total** | **% normal** | **% polyspermic** |
| **M5F5** | **53.51 ± 9.54** | **22.97 ± 10.06** | **23.50 ± 7.15** |
| **M5F20** | **57.59 ± 9.17** | **25.94 ± 8.77** | **16.45 ± 4.93** |
| **M20F5** | **49.23 ± 8.30** | **33.62 ± 11.80** | **17.14 ± 4.78** |
| **M20F20** | **51.61 ± 4.60** | **36.32 ± 4.95** | **12.06 ± 4.83** |

In vitro Maturation and Fertilization under 5 or 20% oxygen tension.
Data presented as mean and ± standard error (p>0.05)

Examining embryo production, our focus was on understanding the effects of high versus low oxygen tension during IVM and IVF, maintaining zygotes in a 5% O2 environment throughout embryonic culture. While the cleavage rate was primarily influenced by IVM (p=0.0003) and IVF (p=0.0022) (Table 2), an interaction emerged between oxygen levels in IVM and IVF regarding embryonic production. Groups exposed to high oxygen levels during maturation and fertilization displayed the highest blastocyst rates. In contrast, those exposed to high oxygen during maturation but low levels during fertilization had the lowest rates (Table 2). Additionally, when assessing intracellular reduced glutathione (GSH) levels in blastocysts, the M20F20 and M5F5 groups exhibited higher GSH levels than the M20F5 and M5F20 groups (p<0.05) (Figure 3A). Differences in IVM oxygen concentrations impacted GSH levels (p=0.0227), contrasting with IVF, where no significant difference was observed (p=0.3801). Furthermore, intracellular ROS levels indicated higher oxidative stress induced by changing oxygen concentrations during procedures (p=0.0016), particularly evident among the M20F20 and M5F5 groups compared to the M20F5 and M5F20 groups (Figure 3B).

**Figure 3.**
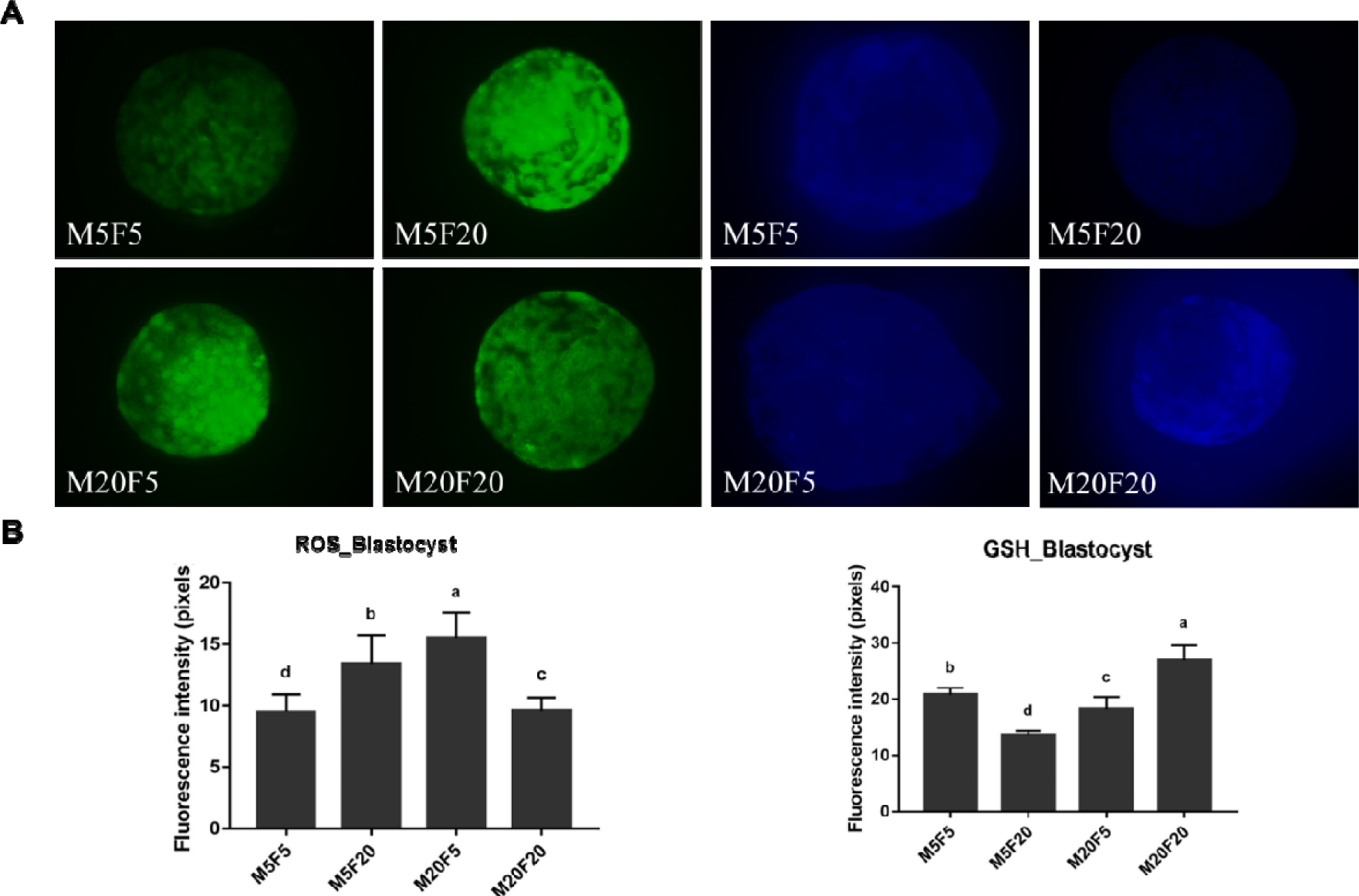
Intracellular levels of ROS and GSH in embryos matured and fertilized under different oxygen tensions (5% and 20%). (A) Epifluorescence photomicrographs of blastocysts stained with CellROX Green for ROS labeling (green) and CellTracker Blue for GSH labeling (blue). All images were taken at 40x magnification, with the same exposure time for fluorescence intensity comparison. (B) Quantification of intracellular levels of reactive oxygen species (ROS) of and reduced glutathione (GSH) in blastocysts matured and fertilized under high and low oxygen tension. The bars in the graph represent the means, and the error bars represent the standard error of the mean. Different letters (a, b, c, d) on the bars indicate a significant difference (p<0.05).

**Table 2:**
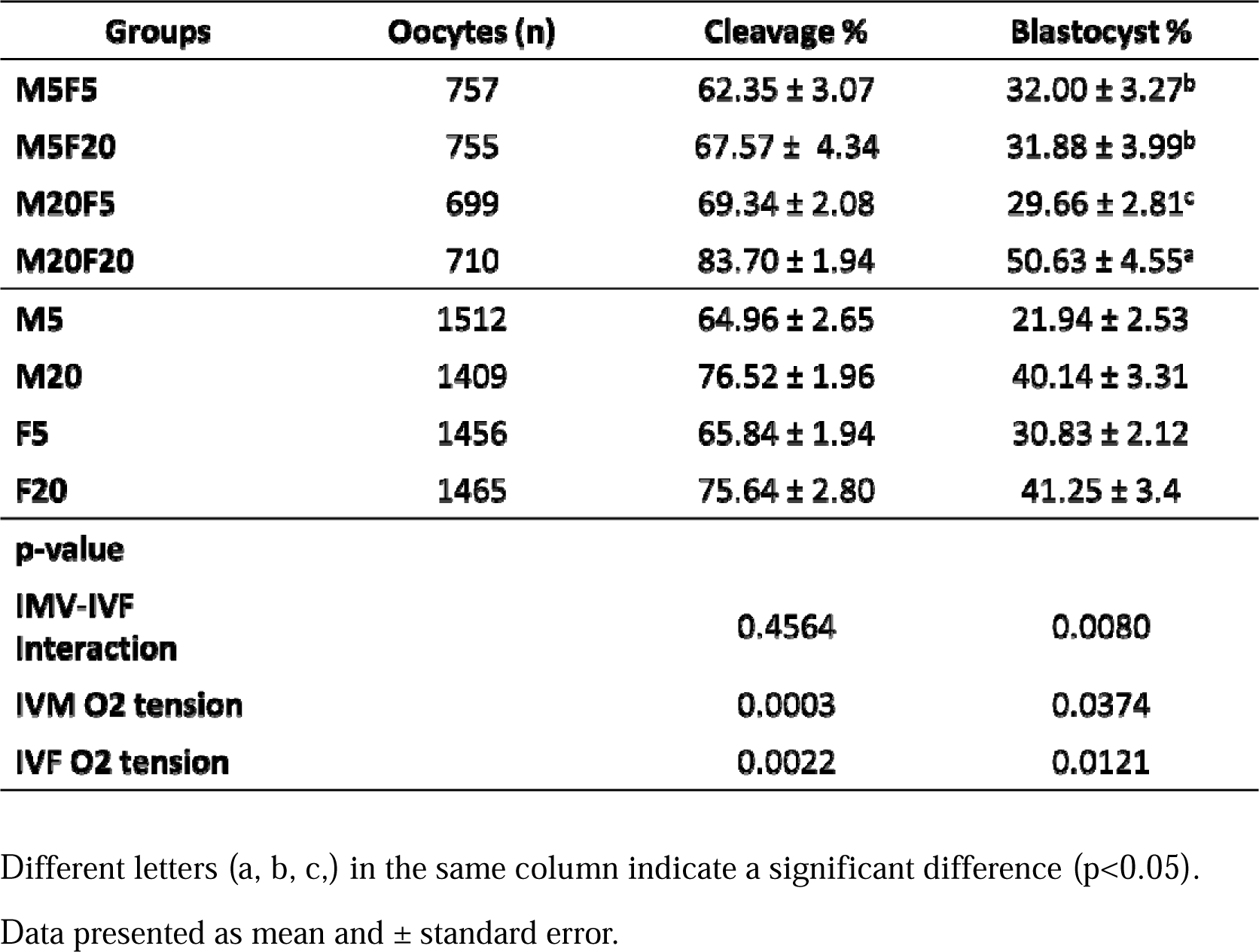
Cleavage and blastocyst rates after in vitro maturation and fertilization at high (20%) or low (5%) oxygen tension.

### Oxygen tension induces changes in epigenetic marks and gene expression in bovine embryos

In this study, we explored epigenetic mark’s responses in embryos subjected to varying oxygen concentrations during maturation and fertilization. Our findings revealed significant impacts of O2 levels on mean fluorescence intensity for H3K9me2 and H3K9me3, with interactions between in vitro maturation (IVM) and in vitro fertilization (IVF) factors (Figure 4A). Notably, H3K9me2 levels were notably higher in embryos matured and fertilized under both low (M5F5) and high (M20F20) oxygen tensions when compared to cross conditions (M5F20 and M20F5) (p < 0.05), mirroring trends seen in H3K9me3 intensities (Figure 4B). Specifically, while H3K9me2 intensities between high and low O2 tension-matured embryos did not differ significantly (p = 0.6495), significant differences were observed in fertilization conditions (p = 0.0125). Similar patterns were found in H3K9me3, indicating significant differences in fertilization conditions (p = 0.0247) but not in maturation conditions (p = 0.5217) (Figure 4B).

**Figure 4.**
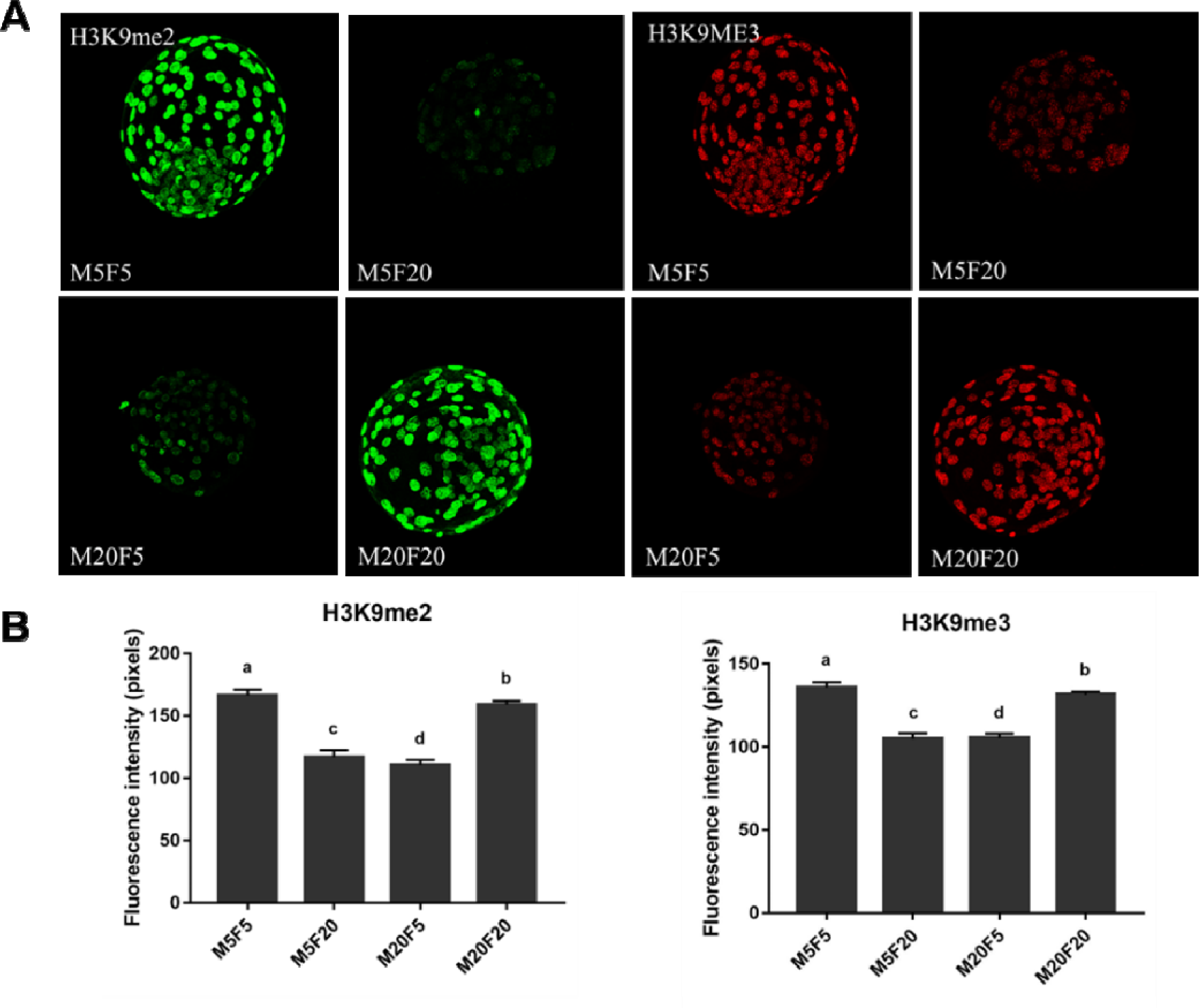
Global levels of H3K9me2 and H3K9me3 in embryos matured and fertilized under different oxygen tensions (5% and 20%). (A) Confocal photomicrographs of blastocysts stained with anti-H3K9me2 antibody (green) and anti-H3K9me3 antibody (red). All images were taken at 40x magnification, with the same exposure time for fluorescence intensity comparison. (B) Quantification of H3K9me2 and H3K9me3 in embryos matured and fertilized under high and low oxygen tension. The bars in the graph represent the means, and the error bars represent the standard error of the mean. Different letters (a, b, c, d) on the bars indicate a significant difference (p< 0.05) by the Kruskal-Wallis test.

Different patterns emerged in the analysis of DNA methylation (5mC) and hydroxymethylation (5hmC) levels. Higher levels of 5mC were evident in embryos matured at low oxygen tension and fertilized either at high tension (M5F20) or at low tension (M5F5), compared to M20F5 and M20F20 groups (p=0.03838) (Figure 5A). When assessing the impact of oxygen tension on in vitro maturation and fertilization, differences in the 5mC mark were observed between embryos matured at low (M5) O2 tension (80566.08±1861.99) and those at 20% O2 (M20) (p<0.05) (Figure 5B). However, 5mC levels in blastocysts fertilized at high (F20) or low (F5) oxygen tension did not significantly differ (p=0.9628) (Figure 5B). Regarding 5hmC levels, differences were noted between blastocysts matured at low (M5) O2 versus those at high (M20) O2 (p=0.0252). Similarly, blastocysts fertilized at low O2 (F5) displayed distinct 5hmC levels (47180.30±2295.35) compared to those fertilized at high O2 (32814.22±1453.39) (p<0.05) (Figure 5B). Ultimately, the qPCR analysis of 17 genes (Supplementary Table) across eight biological replicates revealed no significant interaction (p>0.05, Supplementary Figure S6), emphasizing the distinct main effects of in vitro maturation (MIV). Notably, under low oxygen tension, five genes exhibited differential expression patterns. Specifically, transcripts associated with oxidative stress, including *NRF2* (p=0.01482), *SOD1* (p=0.0410), and *SOD2* (p=0.01374), alongside genes involved in epigenetic remodeling such as *KDM5A* (p=0.04879) and *TET1* (p=0.02798), displayed altered expression profiles in these blastocysts (Supplementary Figure S6).

**Figure 5.**
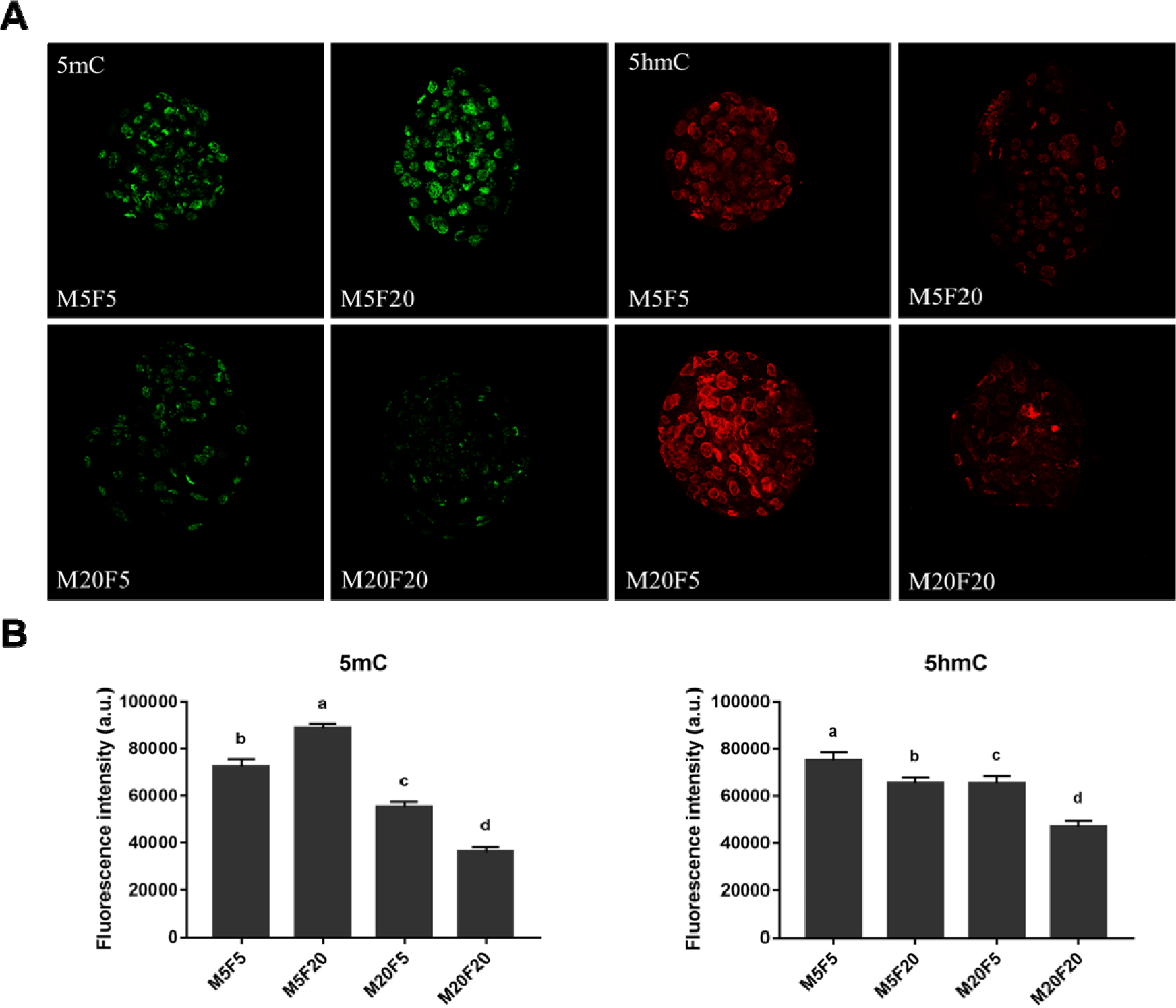
Global levels of 5-mC and 5-hmC in embryos matured and fertilized under different oxygen tensions (5% and 20%). (A) Confocal photomicrographs of blastocysts stained with anti-5-mC antibody (green) and anti-5hmC antibody (red). All images were taken at 40x magnification, with the same exposure time for fluorescence intensity comparison. (B) Quantification of 5mC and 5hmC in embryos matured and fertilized under high and low oxygen tension. The bars in the graph represent the means, and the error bars represent the standard error of the mean. Different letters (a, b, c, d) on the bars indicate a significant difference (p< 0.05) by the Kruskal-Wallis test.

## DISCUSSION

In fibroblasts, the differential expression of HIF2a under high O2 concentrations (20% O2), despite shared regulatory mechanisms with HIF1a, might stem from early cell culture stages, reflecting diverse cellular responses across passages, as noted by Skiles et al. (Skiles et al., 2018). The utilization of 5% oxygen tension in this study might not entirely trigger the hypoxia response cascade, considering that in vivo tissue oxygen concentrations can be lower than those in vitro, aligning with findings from Harvey et al. (A. J. Harvey et al., 2007) and Burr et al. (Burr et al., 2018). H3K9me2 levels were higher under 5% oxygen tension in the fifth passage. This is consistent with the hypoxia-induced histone modifications observed by Chakraborty et al. (Chakraborty et al., 2019). Their study showed that hypoxia can increase global H3K9me2 in mammalian cell lines by activating EHMT2, potentially explaining our findings.

Our study explored the oxygen’s impact on in vitro maturation, revealing that high O2 increased ROS and GSH levels in oocytes, aligning with Hashimoto et al. (2000)(Hashimoto et al., 2000) and Hashimoto (2009)(Hashimoto, 2009). Previous studies also highlighted the intricated interplays between different oxygen levels in in vitro maturation dynamics (del Collado et al., 2017; Kind et al., 2015; Marques et al., 2012). Elevated intracellular GSH in vitro supports embryo development, which is crucial given the increased susceptibility to oxidative damage (Agarwal, Krajcir, Chowdary, & Gupta, 2008). Excessive ROS levels have been shown to cause harmful effects (Al-Gubory, Bolifraud, & Garrel, 2008), while higher GSH concentrations are positively associated with embryo production, notably under high oxygen tension during in vitro maturation.

In a previous report, cumulus cells under high oxygen tension exhibited increased SOD1 expression due to oxidative stress from ROS, compensated by higher intracellular GSH levels (M. B. Harvey, Arcellana-Panlilio, Zhang, Schultz, & Watson, 1995). Low O2 during oocyte in vitro maturation altered genes linked to competence and glucose metabolism (Bermejo-Álvarez, Lonergan, Rizos, & Gutiérrez-Adan, 2010). Elevated oxygen levels increased ROS and oxidative stress-related gene transcription in bovine embryos (Arias, Sanchez, & Felmer, 2012; Corrêa, Rumpf, Mundim, Franco, & Dode, 2008; Rinaudo, Giritharan, Talbi, Dobson, & Schultz, 2006; Wrenzycki et al., 2001). In evaluating oocytes’ in vitro maturation, no significant difference in metaphase II analysis was observed between those matured under high and low oxygen tension. Elevating oxygen tension increased oocytes reaching metaphase II and reduced delayed maturation (Mingoti, Caiado Castro, Méo, Barretto, & Garcia, 2009). Adam et al. (2004) showed that low oxygen levels during in vitro mouse maturation did not impact oocyte development(Abdel, Adam, Takahashi, Katagiri, & Nagano, 2004).

We analyzed the effect of oxygen tension on in vitro fertilization, assessing the fertilization rate of zygotes exposed to high or low O2 tension during IVM and IVF. It was observed that zygotes exposed to low O2 tension during IVF did not result in a higher polyspermy rate. Unlike Hashimoto (2000)(Hashimoto et al., 2000), IVM and IVF under high oxygen tension show a higher polyspermy when 20 mM of glucose is added to the SOFaa medium during in vitro maturation. The importance of low oxygen tension during in vitro fertilization has been discussed in various studies, indicating potential benefits (Arias et al., 2012; Bontekoe et al., 2012; Galli et al., 2001; Takahashi & Kanagawa, 1998; Wale & Gardner, 2016; Yuan et al., 2003) or drawbacks (Bermejo-Álvarez et al., 2010; Pinyopummintr & Bavister, 1995). The consequences of modifications made in IVF raise important yet unresolved questions.

Our results showed that high oxygen levels during in vitro maturation and fertilization improved embryo development rates. This aligns with studies that have reported higher cleavage and blastocyst rates in oocytes exposed to high oxygen environments (de Castro e Paula & Hansen, 2007; Pinyopummintr & Bavister, 1995; Watson et al., 2000), while others have found higher rates at 5% O2 (Bermejo-Álvarez et al., 2010; Hashimoto, 2009; Hashimoto et al., 2000; Kruip, Bevers, & Kemp, 2000; Miller & Rorie, 2000) suggesting that variations in O2 levels during cultivation may account for this discrepancy. Our research revealed that embryos had lower success rates and higher cellular stress when exposed to low oxygen levels during in vitro maturation, as evidenced by increased activity in oxidative stress response pathways such as *SOD1*, *SOD2*, and *NFR2*. This contrasts a previous study by Bermejo-Álvarez et al. (2010), which observed the potential benefits of low oxygen tension during in vitro maturation on bovine oocyte developmental rates and gene expression related to oocyte competence and glucose metabolism(Bermejo-Álvarez et al., 2010). The transcription factor *NRF2*, which activates an important antioxidant pathway, has been linked to the survival and enhanced developmental competence of embryos cultivated under oxidative stress conditions (Amin et al., 2014; del Collado et al., 2017). Unlike Corrêa’s 2009 findings, this study discovered that bovine embryos matured and fertilized under low oxygen tension had lower ROS levels and higher transcript levels of *SOD1* and *SOD2* genes(Corrêa et al., 2008). Further analysis is needed to determine if these embryos experience increased or decreased cellular apoptosis. Future studies should focus on assessing the need for adaptations in the maturation and in vitro fertilization media to better support embryonic development, given the direct link between these environmental changes and metabolism(de Lima et al., 2020).

Furthermore, gene expression analysis revealed elevated levels of epigenetic remodelers *KDM5A* and *TET1* in embryos cultured at lower oxygen tension during in vitro maturation. With their authentic dioxygenase activity, histone demethylases empower chromatin as an oxygen sensor, actively orchestrating the cellular response to hypoxia (Melvin & Rocha, 2012). Our study suggests that the heightened expression of *KDM5A* transcripts in embryos matured under low oxygen tension may arise from its sensitivity to hypoxia, potentially independent of HIF (Batie et al., 2019). Regarding *TET1*, its induction by hypoxia leads to an increase in global 5hmC, offering a potential explanation for the elevated levels of this mark in embryos fertilized under low O2 tension.

In both animal and human contexts, in vitro fertilization represents a crucial shift in the developmental environment. This phase orchestrates significant epigenetic reprogramming, shaping the embryo’s fate (Ross & Sampaio, 2018). Recent attention has focused on the gas atmosphere’s role in influencing the epigenome, particularly the pivotal control of fundamental metabolic processes for embryonic development through O2 concentration (A. J. Harvey et al., 2007). Our study exploring oxygen’s influence on H3K9me2 and H3K9me3 marks in bovine embryos demonstrated elevated levels when maintaining consistent O2 levels during IVM and IVF. These findings aligned with the GSH levels in the embryos. Conversely, embryos matured in one O2 concentration and fertilized in another exhibited heightened ROS levels, indicating that such independent concentration variations might exacerbate oxidative stress that subsequently suppresses marks like H3K9me2/3 (Kietzmann, Petry, Shvetsova, Gerhold, & Görlach, 2017).

Remarkably, embryos matured under low O2 tension demonstrated heightened DNA methylation levels, while 5hmC showed increased levels in low-O2 IVF. Bomfim’s study indicated that embryos cultured under 5% O2 display lower levels of 5mC. However, inferring from this would coincide with our M20F20 group, as we cultured the zygotes in lower O2 concentrations across all groups, rendering it inconclusive. Bennemann et al. 2018 observed higher relative DNA methylation levels in parental pronuclei from oocytes matured in vitro under 5% O2 compared to those at 20%(Bennemann, Grothmann, & Wrenzycki, 2018), suggesting an impact on PGC7/STELLA activity safeguarding TET3-mediated demethylation (Bakhtari & Ross, 2014). TET proteins, involved in oxidizing 5mC to 5hmC during embryo development (Iqbal, Jin, Pfeifer, & Szabó, 2011; Wossidlo et al., 2011), contribute to the heightened 5hmC levels observed in M5F5; however, even in M20F5, there was an increase, hinting that low O2 during fertilization likely amplifies this impact. Those results may have some relationship with sperm oxidation during in vitro fertilization since the paternal genome has to be rapidly demethylated after IVF and ICSI (Abdalla, Yoshizawa, & Hochi, 2009). This group also displayed the lowest embryo developmental rate, likely attributed to an inefficient demethylation process occurring during the development of suboptimal embryos. This inadequacy is recognized as a primary contributor to epigenetic errors in the nuclear reprogramming of clones, sustaining hypermethylation in both DNA and histones (Dean et al., 2001; Santos et al., 2003). In our previous work on cloned embryos, we demonstrated their hypermethylation and the interconnectedness of these marks, highlighting that specific mark modulation can instigate further changes in other marks (Sampaio et al., 2020).

## CONCLUSION

This study aimed to investigate the effect of oxygen tension on in vitro systems, including cell culture, in vitro maturation, and fertilization, and their impact on metabolic processes and epigenetic dynamics. The results showed that fibroblasts cultured under 5% O2 had higher levels of H3K9me2 than those cultured under 20% O2 at initial passages. Then, those embryos matured and fertilized in a high-oxygen environment exhibited enhanced developmental rates. In contrast, embryos matured under low oxygen tension displayed heightened cellular stress, suggesting a multifaceted relationship between oxygen levels and developmental outcomes. Notably, these embryos showcased distinctive epigenetic patterns, featuring reduced DNA methylation and altered 5hmC levels, potentially influenced by oxygen’s impact on DNA demethylation processes during development. Remarkably, variations in O2 tension during IVM and IVF were associated with higher ROS levels and decreased levels of H2K9me2 and H3K9me3, while hypoxia appeared to influence H3K9me2 in somatic cells in early passages. Additionally, the study highlighted differential gene expression related to oxidative stress response pathways under varying oxygen tensions, offering insights into the role of antioxidant mechanisms in ensuring embryo viability. This study emphasizes the need for a nuanced approach to oxygen modulation in assisted reproduction, recognizing its profound effect on embryo development and epigenetic landscapes.

## MATERIALS AND METHODS

### Primary Isolation and Cultivation of Bovine Fibroblasts

Seven skin biopsies (1-2 cm²) were obtained from adult bovine animals at a local, regulated slaughterhouse. Initially, the separation of skin, hide, and fat from the desired tissue was performed using scalpel blades. The sample was washed with saline, antibiotic, and antifungal solution at each step. This was followed by mechanical digestion, achieved by maceration with a scalpel blade, and subsequent enzymatic digestion with 0.1% collagenase incubated at 38.5°C for 3 hours. The cell suspension from each sample was centrifuged at 300 x g for 5 minutes, and the cell pellet was resuspended in Alpha Minimum Essential Medium (α-MEM; GIBCO BRL, Grand Island, NY, USA) supplemented with 15% (v/v) Fetal Bovine Serum (FBS) and 1% antibiotic + antifungal (primary isolation culture medium). The cell suspension from each sample was divided into equal amounts and cultured in 25 cm² tissue culture flasks, with one group incubated under low oxygen tension (5% O2) (Hera cellVios 160i - CO2 incubator, ThermoScientific) and the other under high oxygen tension (20% O2) (Forma Series II incubator, Water-Jacketed CO2 Incubator, ThermoScientific). Cells were cultured until they reached approximately 70% confluence, with medium changes every two days. After this period, cells were passaged using Tryple Express (Invitrogen, Carlsbad, CA, USA) and resuspended in Alpha Minimum Essential Medium (α-MEM; GIBCO BRL, Grand Island, NY, USA) supplemented with 15% (v/v) Fetal Bovine Serum (FBS) and 1% antibiotic + antifungal, followed by replating the cell groups. From the third passage onward, the culture medium was changed to Alpha Minimum Essential Medium (α-MEM; GIBCO BRL, Grand Island, NY, USA) supplemented with 10% (v/v) Fetal Bovine Serum (FBS) and 1% antibiotic (standard culture medium). At each cell passage, the groups resuspended in α-MEM supplemented with 10% FBS and 1% antibiotic were divided into two parts. One part was plated under the same cultural conditions for the subsequent passage. In contrast, the other part was frozen for cell storage in cryotubes using dimethyl sulfoxide (DMSO) and stored in liquid nitrogen. On the fifth, tenth, and twentieth passages, the cell groups resuspended in α-MEM supplemented with 10% FBS and 1% antibiotic were divided into three parts, with two parts maintained as previously described and the third part collected for molecular analysis. After resuspension, the third part of the cells was transferred to an Eppendorf tube, washed in Phosphate-Buffered Saline (PBS), and centrifuged at 300 x g for 5 minutes (repeated three times). Finally, the pellet was snap-frozen by directly immersing the tubes in liquid nitrogen.

### Immunofluorescence for Detection of H3K9me2 in somatic cells

An immunofluorescence assay was conducted on two lines of bovine fibroblasts to identify the H3K9me2 mark. Cells were cultivated on previously sterilized coverslips within 35-mm culture plates (Corning). Cultivation was conducted under high (20%) and low (5%) oxygen tension up to the tenth passage, extending cultures until the twenty-second passage and performing immunofluorescence at these two passages. Following respective cultures, fibroblasts were washed and fixed in 4% paraformaldehyde for 10 minutes, then rinsed thrice in PBS to avoid over-fixation of proteins; all steps described here were conducted on a shaker. Subsequently, the cell membrane was permeabilized for 10 minutes in PBS + 1% TRITON X-100, followed by the addition of blocking buffer, consisting of PBS + 0.3% TRITON X-100 with 1% bovine serum albumin (BSA) (Sigma, St. Louis, MO) for 30 minutes. After the blocking step, cells were incubated overnight at 4°C in a primary antibody solution consisting of a blocking buffer, a mouse antibody anti-H3K9me2—Abcam (ab1220, 1:300), overnight at 4 °C. Cells incubated without primary antibody were used as negative controls for all assays. On the next day, after washing 3× for 10 min each, cells were incubated with secondary antibody Alexa Fluor 488-conjugate goat anti-mouse IgG (Life Tech, cat. #: A-11029) at RT for 1 h. Subsequently, there were three washes of 10 minutes each with washing buffer, with the addition of HOESCHT 33342 (Sigma, St. Louis, MO) in the second wash, diluted for nuclear staining. The coverslips were carefully removed from the plate using a 40:12 needle, mounted on slides containing Prolong Gold Antifade Mountant (Life Tech, cat. # P36935), and examined under a fluorescence microscope (Nikon Eclipse TE 300, Nikon Instruments Inc., Tokyo, Japan) at 200x magnification. Statistical analysis for immunofluorescence involved utilizing the nuclear area obtained with the publicly available ImageJ software, subtracting the dark field value from each nucleus in every photo to remove nonspecific brightness arising from the technique. GraphPad Prism 6 was employed to quantify cellular groups, applying the student’s t-test (p < 0.05) to test for increased or decreased epigenetic marks, generating graphs that facilitate result comprehension.

### In Vitro Embryo Production (IVEP)

The procedures for the maturation, fertilization, and in vitro culture of bovine oocytes/embryos were based on Bomfim et al.’s work (2017) with minor modifications. Ovaries were obtained from slaughterhouses located in Sertãozinho (Barra Mansa), Ipuã (Olhos D’Água), and São João da Boa Vista (Vale do Prata), packed in thermal containers, and immediately transported to the laboratory at a temperature between 33 and 36°C. The aspirated follicular fluid was placed in 50 mL tubes, allowed to settle for oocyte sedimentation, and later observed under a stereomicroscope for COC selection. Following the selection and classification of COCs into grades I, II, or III, a total of 20-25 oocytes were placed in 100 µL drops of maturation medium (B199 tissue culture medium (GIBCO BRL) with Earle’s salts, supplemented with 10% FBS, sodium pyruvate (22 µg/mL), gentamicin (50 µg/mL), 0.5 µg/mL follicle-stimulating hormone (FSH), LH (50µg/mL)), covered with sterile mineral oil (Dow Corning Co., Midland, MI, USA). Subsequently, the oocytes were randomly divided into two groups, matured under an atmosphere of 20% O2 (group M20) or 5% O2, 5% CO2, and 90% N2 (group M5) at a temperature of 38.5°C, for approximately 22 hours, concluding the oocyte maturation process. After maturation, 10 COCs (per group and repetition) were denuded through successive pipetting, and then, oocytes and cumulus cells were separately frozen and stored at −80°C for subsequent molecular analysis. Pools of 10 oocytes from the M5 and M20 groups were stained with the CellROX Green and CellTracker Blue probes to quantify intracellular levels of Reactive Oxygen Species (ROS) and Glutathione (GSH), respectively.

Matured oocytes underwent in vitro fertilization (IVF) with frozen semen from a single Nelore bull (CRV Lagoa, Sertãozinho, SP, Brazil). Capacitated spermatozoa were obtained after separation by Percoll gradient (45% and 90%). The fertilization medium comprised Tyrode’s-lactate stock, 50 µg/ml gentamicin, 22 µg/ml sodium pyruvate, 40 µl/ml PHE (2mM D-penicillamine, 1mM hypotaurine, and 245 µM epinephrine), 5 U/mL heparin, and 6 mg/mL BSA. Each maturation group was subdivided into two groups for IVF under 5% CO2 in air (group F20) or 5% CO2, 5% O2, and 90% N2 (group F5) with maximum humidity for 20 hours at 38.5°C, resulting in four groups (M20F20, M20F5, M5F20, M5F5). The presumptive zygotes were washed and transferred to culture plates (SOFaa medium supplemented with 50 µg/mL gentamicin, 22 µg/mL sodium pyruvate, 5 mg/mL BSA, and 2.5% FBS) under mineral oil. All group developments were carried out in an incubator with low O2 tension (5% CO2, 5% O2, and 90% N2) at 38.5°C, remaining for seven days until embryos reached the blastocyst stage. At this culture period, 96 hours after fertilization (D4), embryo pipetting was performed, and the cleavage rate was visually analyzed with a stereomicroscope at the feeding moment. The embryonic development rate was assessed seven days after fertilization, coinciding with the collection on day 7 of the culture. Blastocysts were collected for ROS, GSH, and epigenetic mark quantification.

### Maturation Rate

To determine the maturation rate (matured oocytes/total oocytes evaluated), a sample of oocytes (15 per repetition) was transferred to plates with 100µL drops of PBS. They were agitated using a micropipette until denuded, fixed for 15 minutes in 4% paraformaldehyde, washed in PBS, and then rinsed in 1mg/mL of PVA in PBS before being stored at 4°C. Oocytes were stained with Hoechst 33342 dye, mounted between a slide and a coverslip in an anti-fading solution (ProLong Gold; Invitrogen, Carlsbad, CA, USA), sealed with nail polish, and observed under an epifluorescence microscope to assess nuclear maturation stage. Oocytes were classified as follows (Patrizio et al., 2007): (1) matured oocytes showing a visible metaphase axis with the extrusion of the first polar body, and (2) immature oocytes lacking a metaphase plate.

### Quantification of Intracellular ROS and GSH Levels

To quantify reactive oxygen species (ROS) formation and reduced glutathione (GSH) in in vitro oocytes and embryos, pools of 10 oocytes matured under high and low oxygen tensions, along with groups of 10 blastocysts, were compared using CellRox Green and CellTracker Blue staining. Analysis was conducted through fluorometric techniques on an epifluorescence microscope using 5µM of CellROX Green (Molecular Probes) and 10 µM of Celltracker Blue (Molecular Probes) for 30 minutes at 37°C. Subsequently, they were fixed in 4% paraformaldehyde for 15 minutes. The oocytes were washed in PBS and placed on slides for epifluorescence analysis. Samples were examined using a 40X objective and a 488 nm wavelength. Images were analyzed by quantifying the fluorescence intensity of the oocytes in grayscale using ImageJ software (NIH; http://rsb.info.nih.gov/ij/), where the software assigns intensity values ranging from 0 to 255 to each pixel.

### Fertilization Rate

The collection procedures for assessing polyspermy rates were based on the work of Alves et al. (2019) with slight modifications. To determine the fertilization rate (total fertilization, normal fertilization, and polyspermy), a sample of zygotes (15 per repetition) was transferred to plates containing 100µL drops of PBS. These were fixed for 15 minutes in 4% paraformaldehyde. Subsequently, they were washed in 1mg/mL PVA solution in PBS and stored at 4°C. The zygotes were stained with Hoechst 33342 dye, mounted between a slide and coverslip in an anti-fade solution (ProLong Gold; Invitrogen, Carlsbad, CA, USA), sealed with enamel, and observed under an epifluorescence microscope.

### Gene expression analysis

The gene expression analysis procedure was conducted following Sampaio’s protocol from 2020, with adaptations tailored to the bovine embryo model. Pools of 10 oocytes and cumulus cells from 5 distinct biological replicates were collected after maturation and blastocysts on day 7 of embryo development for analyzing gene expression patterns associated with oxidative stress and epigenetic remodeling. Total RNA extraction utilized Trizol reagent (Invitrogen, Carlsbad, CA, USA) as per the manufacturer’s instructions, where 1000μL of Trizol was added to each tube. Following the manufacturer’s recommendation, the extracted RNA was eluted in 10 μL of DNase I solution (Invitrogen) plus 1 unit/μL of RNase OUT to degrade contaminating DNA. Individual sample quantification using the NanoDrop Spectrophotometer determined RNA concentrations. Subsequently, RNA was promptly converted to cDNA using the High-Capacity RNA-to-cDNA kit (Life Tech cat # 4387406) following the manufacturer’s protocol and stored at −20 °C until further use. Target genes of interest were those involved in oxidative stress and epigenetic remodeling (Table 1). Relative quantification of mRNA transcripts occurred in 15-μL reactions containing 0.2 μM of previously described primer oligonucleotides, 1x SYBR Green PCR Master Mix (Applied Biosystems), and 2 μL of cDNA diluted 8 times in ultrapure water. All samples were triplicated in the same PCR plate for each gene. Cycling conditions involved initial denaturation at 95°C for 10 minutes, followed by 40 cycles of 95°C for 15 seconds and 60°C for 1 minute. SYBR Green fluorescence was read at the end of each extension step (60°C). Pilot experiments were performed using five different cDNA concentrations ranging from 4-fold to 64-fold dilutions to optimize experimental conditions. PCR product specificity was confirmed through melting curve analysis. The quantity of target transcripts was determined using the formula: E(target)-ΔCt(target)/E(ref)-ΔCt(ref), where E represents the amplification efficiency, and ΔCt signifies the threshold cycle (Ct) difference between control and treated samples. Ct values were obtained after averaging triplicates, while E denotes the estimated mean efficiency for each primer pair (ranging from 90% to 100%) using LinRegPCR software.

### Immunofluorescence for Detection of Epigenetic Marks 5mC, 5hmC, H3K9me2, and H3K9me3 in embryos

Following the protocol outlined by Sampaio et al., 2020, immunostaining was performed on embryos produced under low and high oxygen tension to evaluate the fluorescence intensity of DNA methylation (5mC and 5hmC) and specific histone marks (H3K9me2 and H3K9me3). Blastocysts were fixed in 4% paraformaldehyde for 15 minutes and grouped in pools of 10 embryos from 6 different routines per group. The embryos underwent permeabilization with 0.5% Triton-added PBS for 10 minutes, followed by a PBS wash. To detect 5-mC and 5-hmC, an additional step of DNA denaturation with 4 N HCl for 12 minutes and neutralization with 100 mM Tris HCl solution for 20 minutes was carried out. Next, samples were blocked with PBS solution containing 1% BSA and 0.3 M glycine for 1 hour. Subsequently, blastocysts were incubated with primary antibodies against anti-H3K9me2, anti-H3K9me3, anti-5mC, and anti-5hmC (Abcam) at a 1:200 dilution overnight at 4°C. After 2 hours of washing in 0.1% Triton X-100, the embryos were incubated with secondary antibodies, Goat anti-MOUSE IgG - AlexaFluor 488 (#11008, Invitrogen - Foster City, CA, USA) and Goat anti-rabbit IgG - Texas Red (#111075144, Jackson ImmunoResearch, West Grove, PA, USA) diluted at 1:400 in PBS for 1 hour at room temperature. A negative control blastocyst for the technique (not exposed to primary antibody incubation) was separated for each group. Subsequently, the blastocysts were washed and mounted on slides with 12µL of anti-fade solution (ProLong Antifade Gold reagent with DAPI; Invitrogen) and covered with coverslips. Finally, the samples were analyzed via confocal microscopy using a Leica SP5 microscope (Leica Microsystems, Wetzlar, Germany) with a 40X oil immersion objective. The samples emitted at 590 to 630 nm with 5 slices per embryo. Image analysis for epigenetic mark quantification utilized fluorescence intensity (f.i.) via ImageJ software (NIH; http://rsb.info.nih.gov/ij/), assigning intensity values between 0 and 255 for each pixel.

### Results Analysis

Statistical analyses were performed using R© 3.0.3. For the in vitro embryo production data, the following procedure was adopted. Upon confirming data homogeneity through Levene’s test and data normality via the Shapiro-Wilks test, analysis of variance (ANOVA) was conducted on the quantitative data to assess the responses of embryo production routines, considering their blocks. Pairwise mean comparisons were performed using the Tukey test, with a significance level of p<0.05. If necessary, variables were transformed according to the BOX-COX method. Even after such transformation, if the data did not exhibit a normal distribution, they were evaluated using the Kruskal-Wallis test followed by Fisher’s least significant difference test, and p-values (adjusted, p-adj) were adjusted using the Benjamini-Hochberg method, with a significance level of p-adj<0.05.

## ACKNOWLEDGMENTS

The authors would like to thank the staff and students at the LMMD for their assistance with the sample collections, laboratory procedures, and discussions. The slaughterhouses Olhos D’água and Vale do Prata for gently provided ovaries for this study. The São Paulo Research Foundation supports Rafael Vilar Sampaio — grant number #2015/08807-6; Jessica Brunhara Cruz —FAPESP, grant number #2018/04638-3. This work was granted by São Paulo Research Foundation—FAPESP, grant number #2012/50533-2, #2013/08135-2, #2014/21034-3, #2014/22887-0 and #2014/50947-7; National Counsel of Technological and Scientific Development (CNPq) grant number 465539/2014-9 and CAPES. The funders had no role in study design, data collection and analysis, publication decisions, or manuscript preparation.

## DATA AVAILABILITY

The datasets generated during and analyzed during the current study are available from the corresponding author upon reasonable request.

## REFERENCES

Abdalla, H., Yoshizawa, Y., & Hochi, S. (2009). Active Demethylation of Paternal Genome in Mammalian Zygotes. Journal of Reproduction and Development, 55(4), 356–360. 10.1262/JRD.20234

Abdel, A., Adam, G., Takahashi, Y., Katagiri, S., & Nagano, M. (2004). Effects of oxygen tension in the gas atmosphere during in vitro maturation, in vitro fertilization and in vitro culture on the efficiency of in vitro production of mouse embryos. In Jpn. J. Vet. Res (Vol. 52).

Agarwal, A., Krajcir, N., Chowdary, H., & Gupta, S. (2008). Female Infertility and Assisted Reproduction: Impact of Oxidative Stress. Current Women’s Health Reviews, 4(1), 9–15. 10.2174/157340408783572105

Al-Gubory, K. H., Bolifraud, P., & Garrel, C. (2008). Regulation of Key Antioxidant Enzymatic Systems in the Sheep Endometrium by Ovarian Steroids. Endocrinology, 149(9), 4428–4434. 10.1210/EN.2008-0187

Amin, A., Gad, A., Salilew-Wondim, D., Prastowo, S., Held, E., Hoelker, M., … Tesfaye, D. (2014). Bovine embryo survival under oxidative-stress conditions is associated with activity of the NRF2-mediated oxidative-stress-response pathway. Molecular Reproduction and Development, 81(6), 497–513. 10.1002/MRD.22316

Arias, M. E., Sanchez, R., & Felmer, R. (2012). Evaluation of different culture systems with low oxygen tension on the development, quality and oxidative stress-related genes of bovine embryos produced in vitro. Zygote (Cambridge, England), 20(3), 209–217. 10.1017/S0967199411000025

Bakhtari, A., & Ross, P. J. (2014). DPPA3 prevents cytosine hydroxymethylation of the maternal pronucleus and is required for normal development in bovine embryos. Epigenetics, 9(9), 1271–1279. 10.4161/EPI.32087

Batie, M., Frost, J., Frost, M., Wilson, J. W., Schofield, P., & Rocha, S. (2019). Hypoxia induces rapid changes to histone methylation and reprograms chromatin. Science, 363(6432), 1222–1226. 10.1126/SCIENCE.AAU5870/SUPPL_FILE/AAU5870_DATASET_1.XLSX

Bennemann, J., Grothmann, H., & Wrenzycki, C. (2018). Reduced oxygen concentration during in vitro oocyte maturation alters global DNA methylation in the maternal pronucleus of subsequent zygotes in cattle. Molecular Reproduction and Development, 85(11), 849–857. 10.1002/MRD.23073

Bermejo-Álvarez, P., Lonergan, P., Rizos, D., & Gutiérrez-Adan, A. (2010). Low oxygen tension during IVM improves bovine oocyte competence and enhances anaerobic glycolysis. Reproductive BioMedicine Online, 20(3), 341–349. 10.1016/j.rbmo.2009.12.006

Bomfim, M. M., Andrade, G. M., del Collado, M., Sangalli, J. R., Fontes, P. K., Nogueira, M. F. G., … Perecin, F. (2017). Antioxidant responses and deregulation of epigenetic writers and erasers link oxidative stress and DNA methylation in bovine blastocysts. Molecular Reproduction and Development, 84(12), 1296–1305. 10.1002/MRD.22929

Bontekoe, S., Mantikou, E., van Wely, M., Seshadri, S., Repping, S., & Mastenbroek, S. (2012). Low oxygen concentrations for embryo culture in assisted reproductive technologies. Cochrane Database of Systematic Reviews, (7). 10.1002/14651858.CD008950.PUB2/INFORMATION/EN

Burr, S., Caldwell, A., Chong, M., Beretta, M., Metcalf, S., Hancock, M., … Brewer, A. C. (2018). Oxygen gradients can determine epigenetic asymmetry and cellular differentiation via differential regulation of Tet activity in embryonic stem cells. Nucleic Acids Research, 46(3), 1210–1226. 10.1093/NAR/GKX1197

Chakraborty, A. A., Laukka, T., Myllykoski, M., Ringel, A. E., Booker, M. A., Tolstorukov, M. Y., … Kaelin, W. G. (2019). Histone demethylase KDM6A directly senses oxygen to control chromatin and cell fate. Science (New York, N.Y.), 363(6432), 1217–1222. 10.1126/SCIENCE.AAW1026

Corrêa, G. A., Rumpf, R., Mundim, T. C. D., Franco, M. M., & Dode, M. A. N. (2008). Oxygen tension during in vitro culture of bovine embryos: effect in production and expression of genes related to oxidative stress. Animal Reproduction Science, 104(2–4), 132–142. 10.1016/J.ANIREPROSCI.2007.02.002

de Castro e Paula, L. A., & Hansen, P. J. (2007). Interactions between oxygen tension and glucose concentration that modulate actions of heat shock on bovine oocytes during in vitro maturation. Theriogenology, 68(5), 763–770. 10.1016/J.THERIOGENOLOGY.2007.06.005

de Lima, C. B., dos Santos, É. C., Ispada, J., Fontes, P. K., Nogueira, M. F. G., dos Santos, C. M. D., & Milazzotto, M. P. (2020). The dynamics between in vitro culture and metabolism: embryonic adaptation to environmental changes. Scientific Reports, 10(1). 10.1038/S41598-020-72221-1

Dean, W., Santos, F., Stojkovic, M., Zakhartchenko, V., Walter, J., Wolf, E., & Reik, W. (2001). Conservation of methylation reprogramming in mammalian development: aberrant reprogramming in cloned embryos. Proceedings of the National Academy of Sciences of the United States of America, 98(24), 13734–13738. 10.1073/pnas.241522698

del Collado, M., Da Silveira, J. C., Oliveira, M. L. F., Alves, B. M. S. M., Simas, R. C., Godoy, A. T., … Perecin, F. (2017). In vitro maturation impacts cumulus-oocyte complex metabolism and stress in cattle. Reproduction (Cambridge, England), 154(6), 881–893. 10.1530/REP-17-0134

Fischer, B., & Bavister, B. D. (1993). Oxygen tension in the oviduct and uterus of rhesus monkeys, hamsters and rabbits. Journal of Reproduction and Fertility, 99(2), 673–679. 10.1530/JRF.0.0990673

Galli, C., Crotti, G., Notari, C., Turini, P., Duchi, R., & Lazzari, G. (2001). Embryo production by ovum pick up from live donors. Theriogenology, 55(6), 1341–1357. 10.1016/S0093-691X(01)00486-1

Gaspar, R. C., Arnold, D. R., Corrêa, C. A. P., da Rocha, C. V., Penteado, J. C. T., del Collado, M., … Lopes, F. L. (2015). Oxygen tension affects histone remodeling of in vitro-produced embryos in a bovine model. Theriogenology, 83(9), 1408–1415. 10.1016/J.THERIOGENOLOGY.2015.01.002

Hancock, R. L., Dunne, K., Walport, L. J., Flashman, E., & Kawamura, A. (2015). Epigenetic regulation by histone demethylases in hypoxia. Epigenomics, 7(5), 791–811. 10.2217/EPI.15.24

Harvey, A. J., Navarrete Santos, A., Kirstein, M., Kind, K. L., Fischer, B., & Thompson, J. G. (2007). Differential expression of oxygen-regulated genes in bovine blastocysts. Molecular Reproduction and Development, 74(3), 290–299. 10.1002/MRD.20617

Harvey, M. B., Arcellana-Panlilio, M. Y., Zhang, X., Schultz, G. A., & Watson, A. J. (1995). Expression of Genes Encoding Antioxidant Enzymes in Preimplantation Mouse and Cow Embryos and Primary Bovine Oviduct Cultures Employed for Embryo Coculture. Biology of Reproduction, 53(3), 532–540. 10.1095/BIOLREPROD53.3.532

Hashimoto, S. (2009). Application of In Vitro Maturation to Assisted Reproductive Technology. Journal of Reproduction and Development, 55(1), 1–10. 10.1262/JRD.20127

Hashimoto, S., Minami, N., Takakura, R., Yamada, M., Imai, H., & Kashima, N. (2000). Low Oxygen Tension During In Vitro Maturation is Beneficial for Supporting the Subsequent Development of Bovine Cumulus±Oocyte Complexes. In MOLECULAR REPRODUCTION AND DEVELOPMENT (Vol. 57).

Iqbal, K., Jin, S. G., Pfeifer, G. P., & Szabó, P. E. (2011). Reprogramming of the paternal genome upon fertilization involves genome-wide oxidation of 5-methylcytosine. Proceedings of the National Academy of Sciences of the United States of America, 108(9), 3642–3647. 10.1073/PNAS.1014033108

Kietzmann, T., Petry, A., Shvetsova, A., Gerhold, J. M., & Görlach, A. (2017). The epigenetic landscape related to reactive oxygen species formation in the cardiovascular system. British Journal of Pharmacology, 174(12), 1533–1554. 10.1111/BPH.13792

Kind, K. L., Tam, K. K. Y., Banwell, K. M., Gauld, A. D., Russell, D. L., MacPherson, A. M., … Thompson, J. G. (2015). Oxygen-regulated gene expression in murine cumulus cells. Reproduction, Fertility and Development, 27(2), 407–418. 10.1071/RD13249

Kruip, T. A. M., Bevers, M. M., & Kemp, B. (2000). Environment of oocyte and embryo determines health of IVP offspring. Theriogenology, 53(2), 611–618. 10.1016/S0093-691X(99)00261-7

Loenarz, C., & Schofield, C. J. (2008). Expanding chemical biology of 2-oxoglutarate oxygenases. Nature Chemical Biology, 4(3), 152–156. 10.1038/NCHEMBIO0308-152

Marques, M. G., de Barros, F. R. O., Goissis, M. D., Cavalcanti, P. V., Viana, C. H. C., Assumpção, M. E. O. D., & Visintin, J. A. (2012). Effect of low oxygen tension atmosphere and maturation media supplementation on nuclear maturation, cortical granules migration and sperm penetration in swine in vitro fertilization. Reproduction in Domestic Animals = Zuchthygiene, 47(3), 491–497. 10.1111/J.1439-0531.2011.01909.X

Marsico, T. V., Silva, M. V., Valente, R. S., Annes, K., Rissi, V. B., Glanzner, W. G., & Sudano, M. J. (2023). Unraveling the Consequences of Oxygen Imbalance on Early Embryo Development: Exploring Mitigation Strategies. Animals 2023, Vol. 13, Page 2171, 13(13), 2171. 10.3390/ANI13132171

Martinez, S., & Hausinger, R. P. (2015). Catalytic Mechanisms of Fe(II)- and 2-Oxoglutarate-dependent Oxygenases. The Journal of Biological Chemistry, 290(34), 20702–20711. 10.1074/JBC.R115.648691

Melvin, A., & Rocha, S. (2012). Chromatin as an oxygen sensor and active player in the hypoxia response. Cellular Signalling, 24(1), 35–43. 10.1016/J.CELLSIG.2011.08.019

Menezo, Y. J. R., Silvestris, E., Dale, B., & Elder, K. (2016). Oxidative stress and alterations in DNA methylation: two sides of the same coin in reproduction. Reproductive Biomedicine Online, 33(6), 668–683. 10.1016/J.RBMO.2016.09.006

Miller, G. F., & Rorie, R. W. (2000). Effect of oxygen concentration during oocyte maturation on subsequent bovine embryo cleavage and development in vitro. Effect of Oxygen Concentration during Oocyte Maturation on Subsequent Bovine Embryo Cleavage and Development in Vitro., (No. 478), 43–44.

Mingoti, G. Z., Caiado Castro, V. S. D., Méo, S. C., Barretto, L. S. S., & Garcia, J. M. (2009). The effect of interaction between macromolecule supplement and oxygen tension on bovine oocytes and embryos cultured in vitro. Zygote (Cambridge, England), 17(4), 321–328. 10.1017/S0967199409005450

Nabenishi, H., Ohta, H., Nishimoto, T., Morita, T., Ashizawa, K., & Tsuzuki, Y. (2012). The effects of cysteine addition during in vitro maturation on the developmental competence, ROS, GSH and apoptosis level of bovine oocytes exposed to heat stress. Zygote (Cambridge, England), 20(3), 249–259. 10.1017/S0967199411000220

Oyamada, T., & Fukui, Y. (2004). Oxygen tension and medium supplements for in vitro maturation of bovine oocytes cultured individually in a chemically defined medium. The Journal of Reproduction and Development, 50(1), 107–117. 10.1262/JRD.50.107

Pinyopummintr, T., & Bavister, B. D. (1995). Optimum gas atmosphere for in vitro maturation and in vitro fertilization of bovine oocytes. Theriogenology, 44(4), 471–477. 10.1016/0093-691X(95)00219-X

Rinaudo, P. F., Giritharan, G., Talbi, S., Dobson, A. T., & Schultz, R. M. (2006). Effects of oxygen tension on gene expression in preimplantation mouse embryos. Fertility and Sterility, 86(4 Suppl), 1265.e1–1265.e36. 10.1016/J.FERTNSTERT.2006.05.017

Ross, P. J., & Sampaio, R. V. (2018). Epigenetic remodeling in preimplantation embryos: cows are not big mice. Animal Reproduction, 15(3), 204–214. 10.21451/1984-3143-AR2018-0068

Sampaio, R. V, Sangalli, J. R., De Bem, T. H. C., Ambrizi, D. R., del Collado, M., Bridi, A., … Meirelles, F. V. (2020). Catalytic inhibition of H3K9me2 writers disturbs epigenetic marks during bovine nuclear reprogramming. Scientific Reports, 10(1).

Santos, F., Zakhartchenko, V., Stojkovic, M., Peters, A., Jenuwein, T., Wolf, E., … Dean, W. (2003). Epigenetic Marking Correlates with Developmental Potential in Cloned Bovine Preimplantation Embryos. Current Biology, 13(13), 1116–1121.

Semenza, G. L. (2002). Signal transduction to hypoxia-inducible factor 1. Biochemical Pharmacology, 64(5–6), 993–998. 10.1016/S0006-2952(02)01168-1

Semenza, G. L. (2003). Targeting HIF-1 for cancer therapy. Nature Reviews. Cancer, 3(10), 721–732. 10.1038/NRC1187

Skiles, W. M., Kester, A., Pryor, J. H., Westhusin, M. E., Golding, M. C., & Long, C. R. (2018). Oxygen-induced alterations in the expression of chromatin modifying enzymes and the transcriptional regulation of imprinted genes. Gene Expression PatternsL: GEP, 28, 1–11. 10.1016/J.GEP.2018.01.001

Takahashi, Y., & Kanagawa, H. (1998). Effect of Oxygen Concentration in the Gas Atmosphere during In Vitro Insemination of Bovine Oocytes on the Subsequent Embryonic Development In Vitro. Journal of Veterinary Medical Science, 60(3), 365–367. 10.1292/JVMS.60.365

van der Knaap, J. A., & Verrijzer, C. P. (2016). Undercover: gene control by metabolites and metabolic enzymes. Genes & Development, 30(21), 2345–2369. 10.1101/GAD.289140.116

Wale, P. L., & Gardner, D. K. (2016). The effects of chemical and physical factors on mammalian embryo culture and their importance for the practice of assisted human reproduction. Human Reproduction Update, 22(1), 2–22. 10.1093/HUMUPD/DMV034

Watson, A. J., De Sousa, P., Caveney, A., Barcroft, L. C., Natale, D., Urquhart, J., & Westhusin, M. E. (2000). Impact of bovine oocyte maturation media on oocyte transcript levels, blastocyst development, cell number, and apoptosis. Biology of Reproduction, 62(2), 355–364. 10.1095/BIOLREPROD62.2.355

Wossidlo, M., Nakamura, T., Lepikhov, K., Marques, C. J., Zakhartchenko, V., Boiani, M., … Walter, J. (2011). 5-Hydroxymethylcytosine in the mammalian zygote is linked with epigenetic reprogramming. Nature Communications, 2(1). 10.1038/NCOMMS1240

Wrenzycki, C., Herrmann, D., Keskintepe, L., Martins, A., Sirisathien, S., Brackett, B., & Niemann, H. (2001). Effects of culture system and protein supplementation on mRNA expression in pre-implantation bovine embryos. Human Reproduction (Oxford, England), 16(5), 893–901. 10.1093/HUMREP/16.5.893

Yuan, Y. Q., Van Soom, A., Coopman, F. O. J., Mintiens, K., Boerjan, M. L., Van Zeveren, A., … Peelman, L. J. (2003). Influence of oxygen tension on apoptosis and hatching in bovine embryos cultured in vitro. Theriogenology, 59(7), 1585–1596. 10.1016/S0093-691X(02)01204-9

