## Supplemental Files for "Artificial environment impact: O2 concentration changes between IVM and IVF alter embryo production, metabolism, and epigenetic marks"

**Supplementary Figure S1**

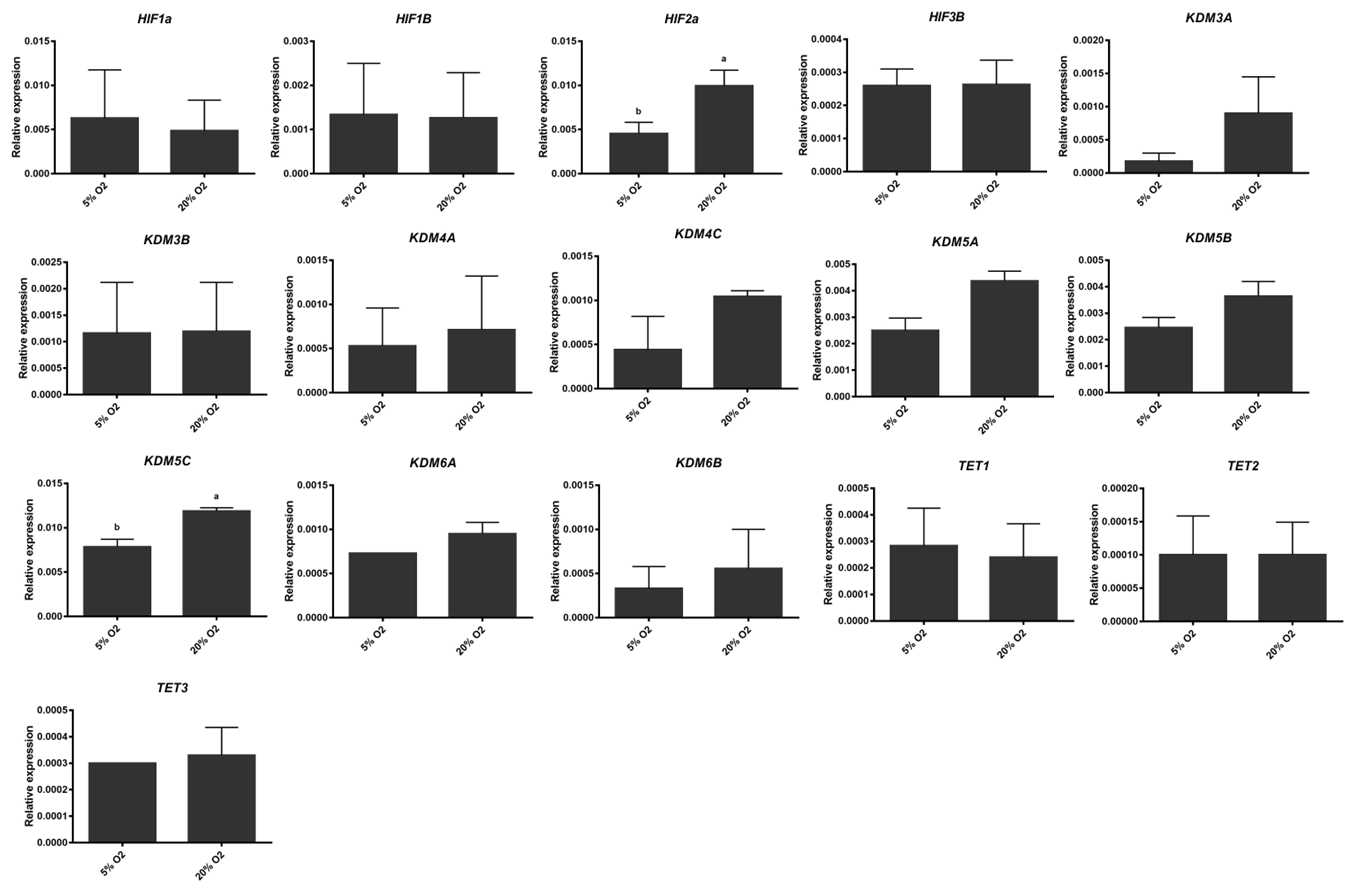

**Supplementary Figure S1**. Relative expression of target genes related to oxidative stress response and epigenetic remodeling differentially expressed bovine fibroblasts cultured at passage 5 under high or low oxygen tension. The bars in the graph represent the means, and the error bars represent the standard error of the mean. Different letters (a, b) above the bars indicate a significant difference (p<0.05).

**Supplementary Figure S2**

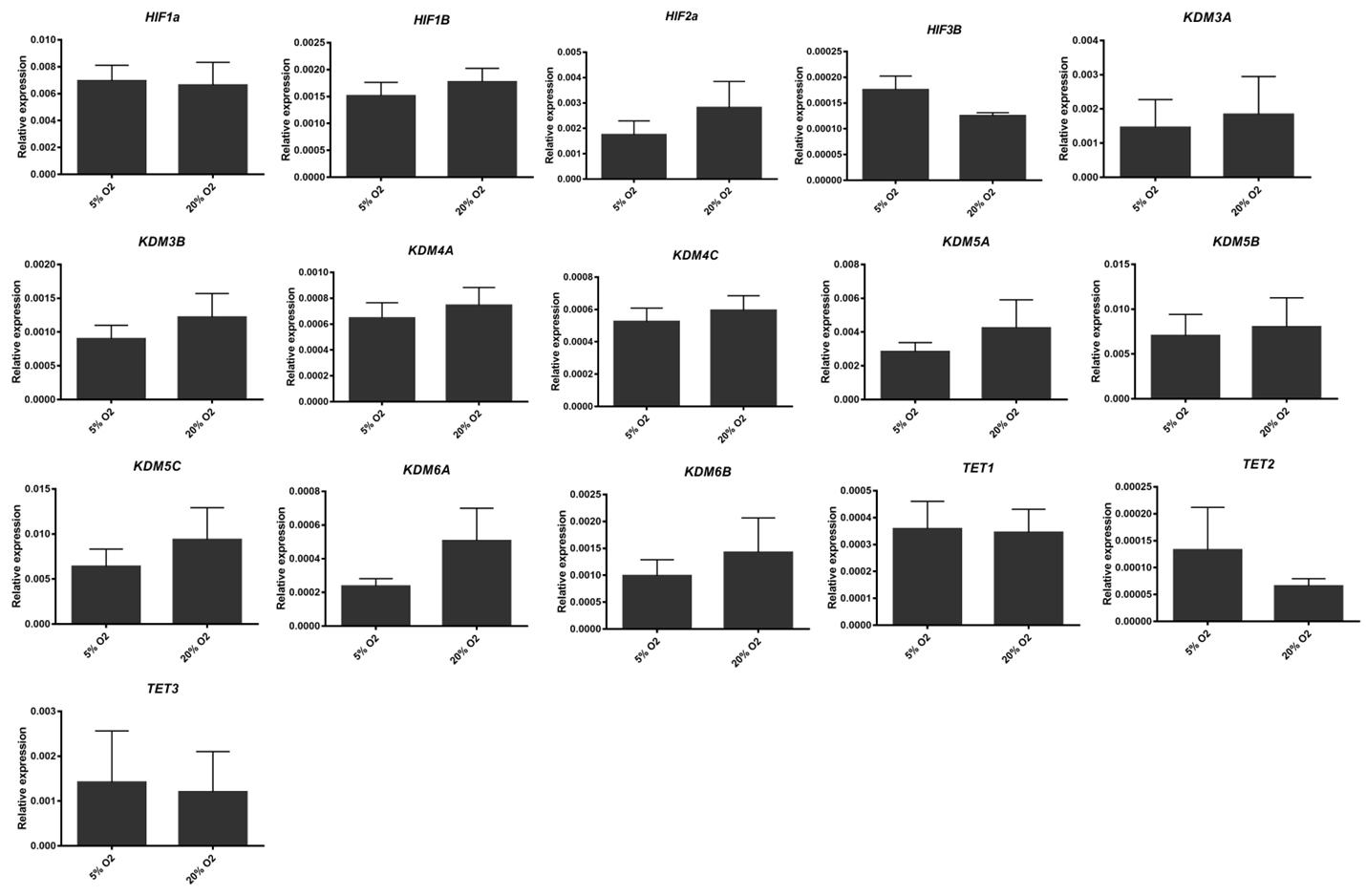
**Supplementary Figure S2**. Relative expression of target genes related to oxidative stress response and epigenetic remodeling differentially expressed bovine fibroblasts cultured at passage 20 under high or low oxygen tension. The bars in the graph represent the means, and the error bars represent the standard error of the mean. No difference was found (p>0.05).

**Supplementary Figure S3**

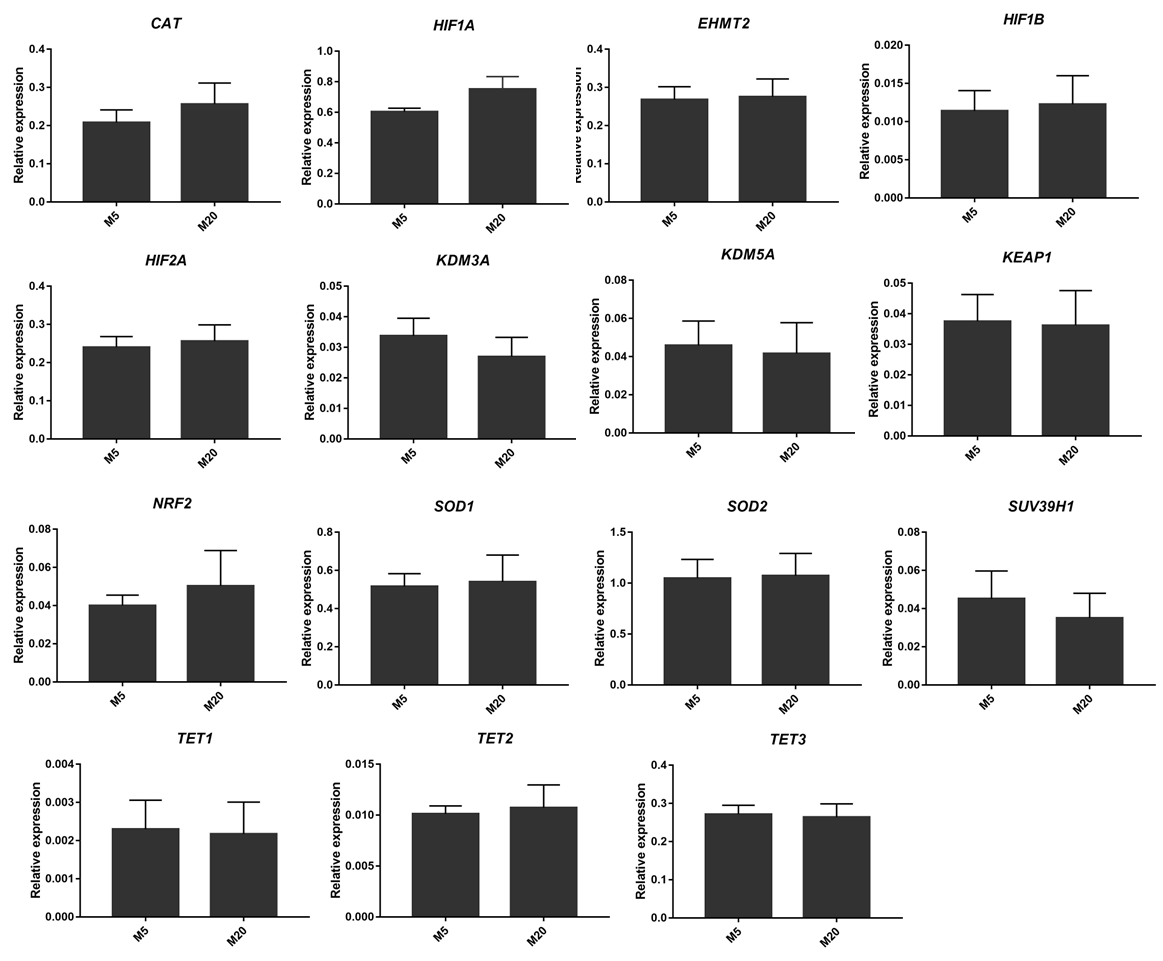
 **Supplementary Figure S3**. Relative expression of target genes related to oxidative stress response and epigenetic remodeling differentially expressed from oocytes matured under high or low oxygen tension. The bars in the graph represent the means, and the error bars represent the standard error of the mean. No significant difference was found (p>0.05).

**Supplementary Figure S4**

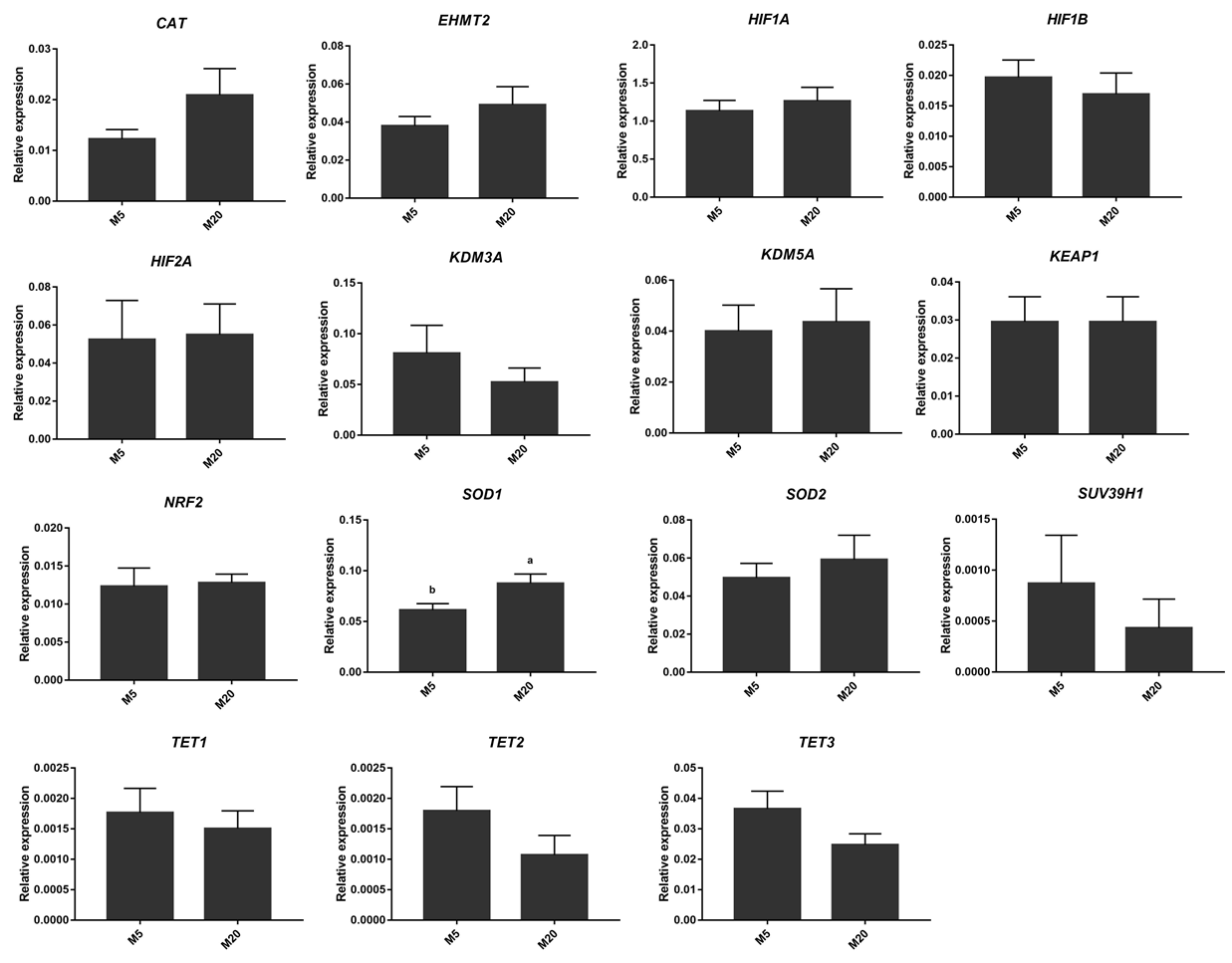
 **Supplementary Figure S4**. Relative expression of target genes related to oxidative stress response and epigenetic remodeling is differently expressed in cumulus cells from oocytes maturated under high or low oxygen tension. The bars in the graph represent the means, and the error bars represent the standard error of the mean. Different letters (a, b) above the bars indicate a significant difference (p<0.05).

**Supplementary Figure S5**

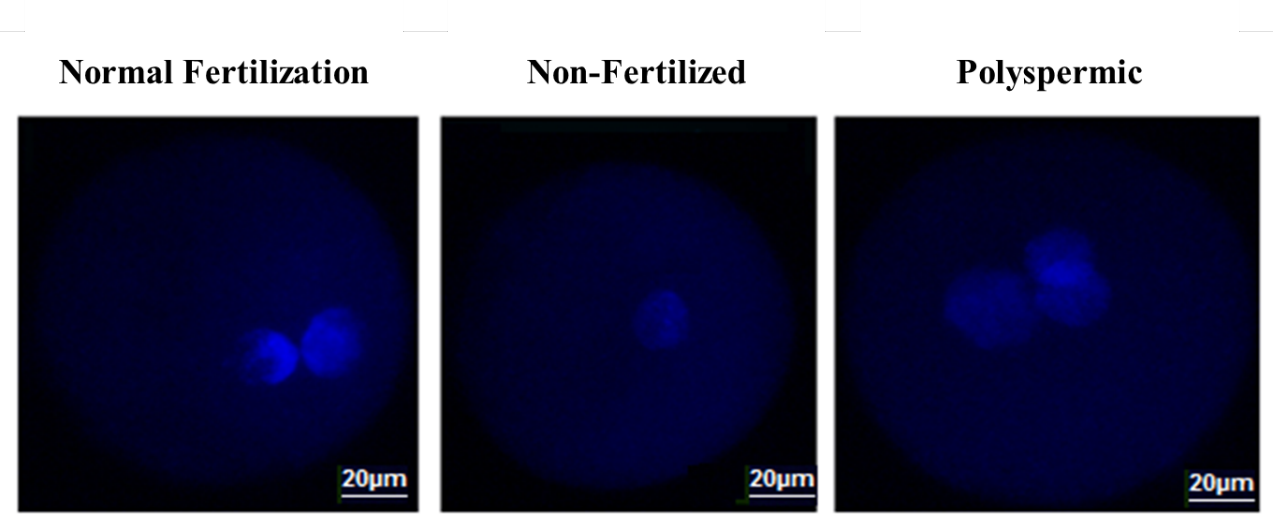
 **Supplementary Figure S5**. Fluorescence photomicrograph of zygotes produced under high and low oxygen tension during in vitro maturation and fertilization, stained with Hoechst 33342, depicting a zygote with normal fertilization, non-fertilized and polyspermic.

**Supplementary Figure S6**

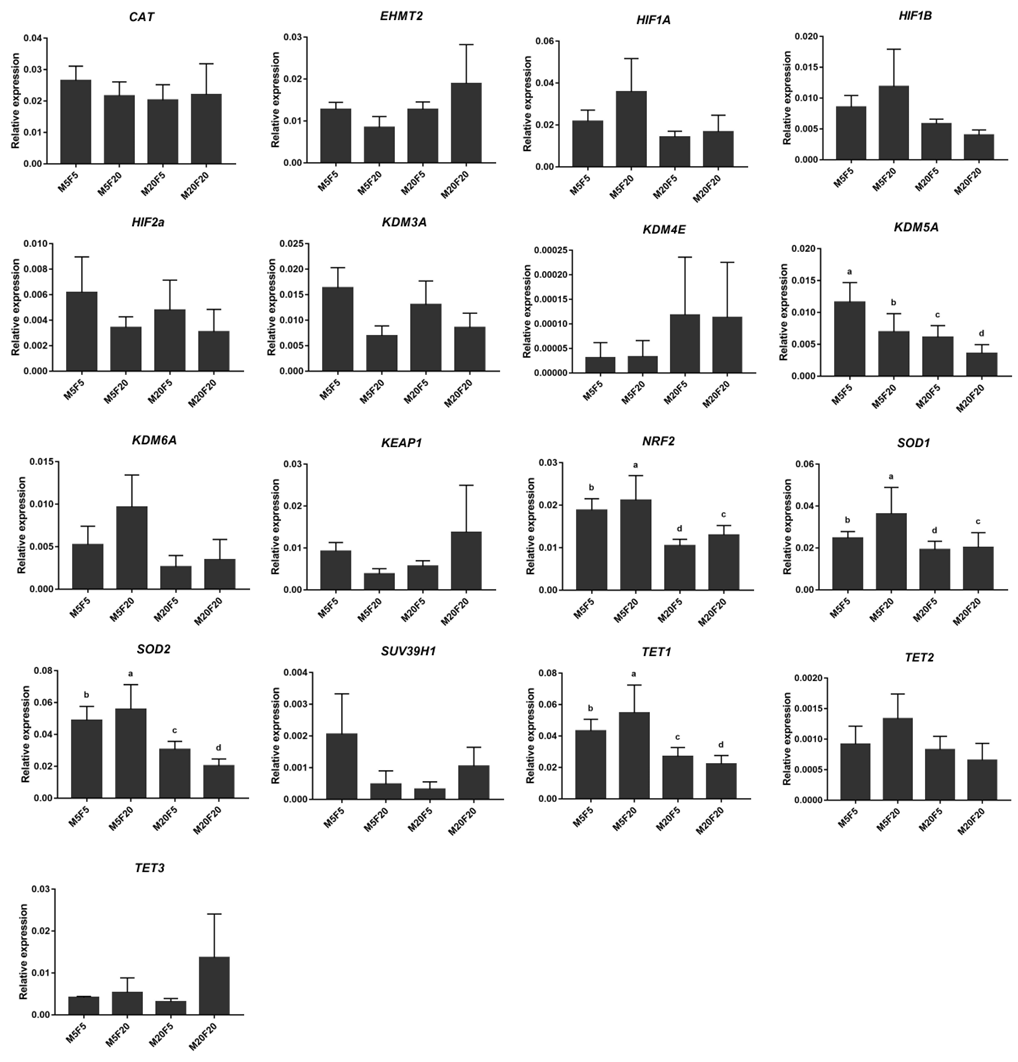

**Supplementary Figure S6**. Relative expression of target genes related to oxidative stress response and epigenetic remodeling in embryos maturated and fertilized under high or low oxygen tension. The bars in the graph represent the means, and the error bars represent the standard error of the mean. Different letters (a, b) above the bars indicate a significant difference (p<0.05).

**Supplementary Table**

| **Gene Symbol** | **Gene Name** | **Sequence (5’-3’)** | **AT (°C)** | **Source** |
| --- | --- | --- | --- | --- |
| *Tet1* | *Ten-eleven translocation methylcytosine dioxygenase 1* | F: TGCCTACTTGCAACTGTCTTGATC | 60 | (R. V Sampaio et al., 2020) |
|  |  | R: TCTATCCTTACTGCATTTCCTTTTTG |  |  |
| *Tet2* | *Ten-eleven translocation methylcytosine dioxygenase 2* | F: GGATACACCTGTCAAGACTCAGTATGA | 60 | (R. V. Sampaio et al., 2020) |
|  |  | R: GGACCTGCTCCTAGATGGGTATAA |  |  |
| *Tet3* | *Ten-eleven translocation methylcytosine dioxygenase 3* | F: TGCGGCCTCAACGATGA | 60 | (R. V Sampaio et al., 2020) |
|  |  | R: GGAACAACCGAAGGAGAAGGA |  |  |
| *KDM3A* | *Lysine (K)-Specific Demethylase 3A* | F: AGTGGCCTGCAATAATGTACAAA | 60 | (R. V. Sampaio, 2016) |
|  |  | R:GAAAGACGAAGATTGTTTACATCC |  |  |
| *KDM3B* | *Lysine (K)-Specific Demethylase 3B* | F: ACTGGTTGGCGGATTTAGCA | 60 | (R. V. Sampaio, 2016) |
|  |  | R: ACGGAGAGACCTTTGCAGTG |  |  |
| *KDM4A* | *Lysine (K)-Specific Demethylase 4A* | F: TGCTTCTGTAGCCTTGACCG | 60 | (R. V. Sampaio, 2016) |
|  |  | R: GGTCAGGCAGAAATCGCAGA |  |  |
| *KDM4C* | *Lysine (K)-Specific Demethylase 4C* | F: CCCAAGCAGTTCTCAAGGGT | 60 | (R. V. Sampaio, 2016) |
|  |  | R: GCTTCCTGGGTGATCTTGTCA |  |  |
| *KDM5A* | *Lysine (K)-Specific Demethylase 5A* | F: GGTGTTTGAGCTTGTGCCTG | 60 | (Glanzner et al., 2018) |
|  |  | R: TGTAACACGACTGACCCACG |  |  |
| *KDM5B* | *Lysine (K)-Specific Demethylase 5B* | F: GACGTGTGCCAGTTTTGGAC | 60 | (Glanzner et al., 2018) |
|  |  | R: TCGAGGACACAGCACCTCTA |  |  |
| *KDM5C* | *Lysine (K)-Specific Demethylase 5C* | F: ATCACTACCCCTGCCTGGAT | 60 | (Glanzner et al., 2018) |
|  |  | R: CTGAGGTCCTGCGCAGATAG |  |  |
| *KDM6A* | *Lysine (K)-Specific Demethylase 6A* | F: TGGAACAGCTCCGTGCAAAT | 60 | (Glanzner et al., 2018) |
|  |  | R: GAGACACGCTAGGCACTCTG |  |  |
| *KDM6B* | *Lysine (K)-Specific Demethylase 6B* | F: GGGAGACTATCAGCGCCTTC | 60 | (Glanzner et al., 2018) |
|  |  | R: AGCGGTACACAGGGATGTTG |  |  |
| *HIF1α* | *HypoxiaInducibleFactor 1α* | F: TGAAGGCACAGATGAATTGCTT | 60 | (Harvey, 2004) |
|  |  | R: GTTCAAACTGAGTTAATCCCATGTATTT |  |  |
| *HIF1B* | *HypoxiaInducibleFactor 1B* | F: ATTAAGCGGCGACCAGGATT | 60 | (Harvey, , 2004) |
|  |  | R: CTCATCATCCGACCTGGCAA |  |  |
| *HIF2α* | *HypoxiaInducibleFactor 2α* | F: GCACCATGTCAAACATCTTCCA | 60 | (Harvey, 2004) |
|  |  | R: TCAAAGAAGGCGAAGGACACA |  |  |
| *HIF3β* | *HypoxiaInducibleFactor3 β* | F: GCACGGCGTTCTTTCTTCTG | 60 | (Harvey et al., 2004) |
|  |  | R: TGCAGAAGCTTTTTCGATCTTTTT |  |  |
| *ACTB* | *Actin beta* | F: CAGCAGATGTGGATCAGCAAGC | 60 | (R. V. Sampaio, 2016) |
|  |  | R: AACGCAGCTAACAGTCCGCC |  |  |
| H2A | Histone H2A | F: GAGGAGCTGAACAAGCTGTTG | 60 | (R. V Sampaio et al., 2020) |
|  |  | R: TTGTGGTGGCTCTCAGTCTTC |  |  |
| *KEAP1* | *Kelch-like ECH-associated protein 1* | F: TCACCAGGGAAGGATCTACG | 60 | (Amin et al., 2014) |
|  |  | R: AGCGGCTCAACAGGTACAGT |  |  |
| *SOD1* | *Superoxide Dismutase 1* | F: AGAGGCATGTTGGAGACAT | 60 | (Amin et al., 2014) |
|  |  | R: CAGCGTTGCCAGTCTTTGTA |  |  |
| *NRF2* | *Nuclear factor erythroid 2-related factor 2* | F: AGCTGCCAAGATGGTGAAAC | 60 | (Amin et al., 2014) |
|  |  | R: ACTCTGCAGCAACATCCTGA |  |  |
| *CAT* | *Catalase* | F: TGGGACCCAACTATCTCCAG | 60 | (Amin et al., 2014) |
|  |  | R: AAGTGGGTCCTGTGTTCCAG |  |  |
| *SOD2* | *Superoxide dismutase-2* | F: AGAGGCAATGTTGGAGACCTG | 60 | (Amin et al., 2014) |
|  |  | R: TCTCGTTGCCAGTCTTTGTA |  |  |

**References**

Amin, A., Gad, A., Salilew-Wondim, D., Prastowo, S., Held, E., Hoelker, M., … Tesfaye, D. (2014). Bovine embryo survival under oxidative-stress conditions is associated with activity of the NRF2-mediated oxidative-stress-response pathway. *Molecular Reproduction and Development*, *81*(6), 497–513. https://doi.org/10.1002/MRD.22316

Glanzner, W. G., Rissi, V. B., de Macedo, M. P., Mujica, L. K. S., Gutierrez, K., Bridi, A., … Bordignon, V. (2018). Histone 3 lysine 4, 9, and 27 demethylases expression profile in fertilized and cloned bovine and porcine embryos. *Biology of Reproduction*, *98*(6), 742–751.

Harvey, a J., Kind, K. L., Pantaleon, M., Armstrong, D. T., & Thompson, J. G. (2004). Oxygen-regulated gene expression in bovine blastocysts. *Biology of Reproduction*, *71*(4), 1108–1119. https://doi.org/10.1095/biolreprod.104.028639

Sampaio, R. V. (2016). *Modificações epigenéticas da cromatina e sua relação com a reprogramação nuclear de bovinos*. https://doi.org/10.11606/T.10.2016.TDE-27082015-114515

Sampaio, R. V., Sangalli, J. R., De Bem, T. H. C., Ambrizi, D. R., del Collado, M., Bridi, A., … Meirelles, F. V. (2020). Catalytic inhibition of H3K9me2 writers disturbs epigenetic marks during bovine nuclear reprogramming. *Scientific Reports*, *10*(1). https://doi.org/10.1038/S41598-020-67733-9
